# Integron Gene Cassettes Harboring Novel Variants of D-Alanine-D-Alanine Ligase Confer High-level Resistance to D-Cycloserine

**DOI:** 10.1101/2020.05.09.085589

**Authors:** Md. Ajijur Rahman, Frank Kaiser, Shirin Jamshidi, Khondaker Miraz Rahman, Peter Mullany, Adam P. Roberts

## Abstract

During a PCR-based screen of antibiotic resistance genes (ARGs) associated with integrons in saliva-derived metagenomic DNA of healthy human volunteers, two novel variants of genes encoding a D-alanine-D-alanine ligase (*ddl6* and *ddl7*) located within gene cassettes in the first position of a reverse integron were identified. *Treponema denticola* was identified as the likely host of the *ddl* cassettes. Both *ddl6* and *ddl7* conferred high level resistance to D-cycloserine when expressed in *Escherichia coli* with *ddl7* conferring four-fold higher resistance to D-cycloserine compared to *ddl6*. A SNP was found to be responsible for this difference in resistance phenotype between both *ddl* variants. Molecular dynamics simulations were used to explain the mechanism of this phenotypic change at the atomic scale. A hypothesis for the evolutionary selection of *ddl* containing integron gene cassettes is proposed, based on molecular docking of plant metabolites within the ATP and D-cycloserine binding pockets of Ddl.

**Significance Statement:** The drivers of antimicrobial resistance are varied and numerous. One such hypothesis is that compounds not normally considered to be antibiotics may be able to select for genes that can provide resistance to conventional antibiotics. Here we provide evidence that dietary flavonoids are likely to provide the selective pressure for two novel variants of the *ddl* gene, encoding D-alanine-D-alanine ligases, to be maintained as the first gene cassette of a reverse integron detectable in the human oral cavities from both the UK and Bangladesh. We show that Ddl is functional, able to confer resistance to D-cycloserine and that the dietary flavonoids quercetin and apigenin are likely able to compete with both ATP and D-cycloserine within their Ddl binding sites.

## Introduction

Globally, the emergence of antibiotic resistance in clinically important pathogens pose a significant threat to human health. Antibiotic resistance genes (ARGs) carried on various mobile genetic elements (MGEs), including plasmids, transposons and integron gene cassettes (GCs) are responsible for the emergence, and worldwide dissemination, of multidrug-resistance. Identifying the principal reservoirs of ARGs (1, 2) as well as understanding the drivers for their evolutionary selection and spread (3) are fundamental to tackling the problem of antibiotic resistance.

Integrons are genetic elements capable of capturing and expressing open reading frames (ORFs) embedded in GCs. They are well known for their role in dissemination of ARGs particularly in gram-negative pathogens. The basic structure of an integron is composed of a gene encoding a DNA integrase (*intI*), an integron associated recombination site; *attI* and a gene cassette promoter (*Pc)*, which expresses the inserted cassettes within the GC array. The integron integrase (IntI) catalyses recombination between the cassette associated recombination site; *attC* and the attachment site; *attI*, within the integron platform (4-6). More than 130 MGE associated ARGs have been detected in integrons, conferring resistance to most classes of antibiotics including aminoglycosides, chloramphenicol, beta-lactams, trimethoprim and streptothricin (7) (8). Mobile integrons, which usually carry ARGs, are associated with MGEs such as transposons or plasmids and possess a few cassettes in the cassette array, whereas chromosomal integrons usually have a large GC array encoding proteins with diverse functions including adaptation to stress, virulence and resistance to xenobiotics. However, the majority of them encode proteins of unknown functions (9). For instance, among the 70 ORFs detected within 47 gene cassettes in the chromosomal integron of *Treponema denticola* (10), an oral pathogen associated with periodontitis, only five ORFs matched with proteins of known functions (last confirmed August 2019).

This article reports the detection of two novel integron-located genes encoding D-alanine-D-alanine ligases (Ddl). Ddls are ATP-dependent enzymes that play a critical role in the synthesis of peptidoglycan during the cytoplasmic stage by catalysing the formation of the D-ala-D-ala dipeptide required for cross-linking peptidoglycan strands during cell wall synthesis (11). The gene encoding Ddl is a house-keeping gene and has always been found in the chromosome, never in a MGE. The formation D-Ala-D-Ala dipeptide catalysed by Ddl is critical for bacterial growth. As there is no human homologue, Ddl is considered an excellent target for designing new antibiotics (12). D-cycloserine is the only inhibitor of Ddl used in the clinic, however, due to the severe neurologic side effects, its use is currently reserved for the treatment of multidrug resistant (MDR) and extensively-drug resistant (XDR) tuberculosis.

We demonstrate D-cycloserine resistance associated with these integron located *ddls* and present an experimentally supported hypothesis as to why *ddl*, a housekeeping gene, has evolved to be on an integron gene cassette.

## Results

### Two novel variants of *ddl* were discovered from the human oral cavity located immediately downstream of the core structure of a reverse integron

During a PCR screen (all primers listed in Table S4) for the presence of integron GCs carrying ARGs in the saliva-derived metagenomic DNA of healthy volunteers from the UK (n=11) and Bangladesh (n=12), two natural variants of *ddl* located within the first GC of a reverse integron were discovered. In the libraries of PCR amplicons of integrons and associated GCs, a total of five clones (three from the UK and two from Bangladesh) were recovered carrying *ddl* predicted to encode a D-alanine-D-alanine ligase. The genetic organisation of the *ddl* inserts in the clones are shown in Fig. 1A. The *intI* and the *ddl* gene cassettes are oriented in the same direction in reverse integrons (where the integrase is orientated in the opposite direction compared to usual integrons) of *T. denticola* (10). Two putative integrase binding sites [L or S2 and R or S1], two direct repeat sequences (DR1 and DR2) and a putative *attI/attC* recombination site were detected (Fig. 1B). The stop codon of the ORF encoding *ddl* was located on the 7 bp *attC* core site, R′′ (Fig. 1C). The R′′ site is followed by the L′′ site -which covers the priming site of the reverse primer MARS2.

**Figure 1.**
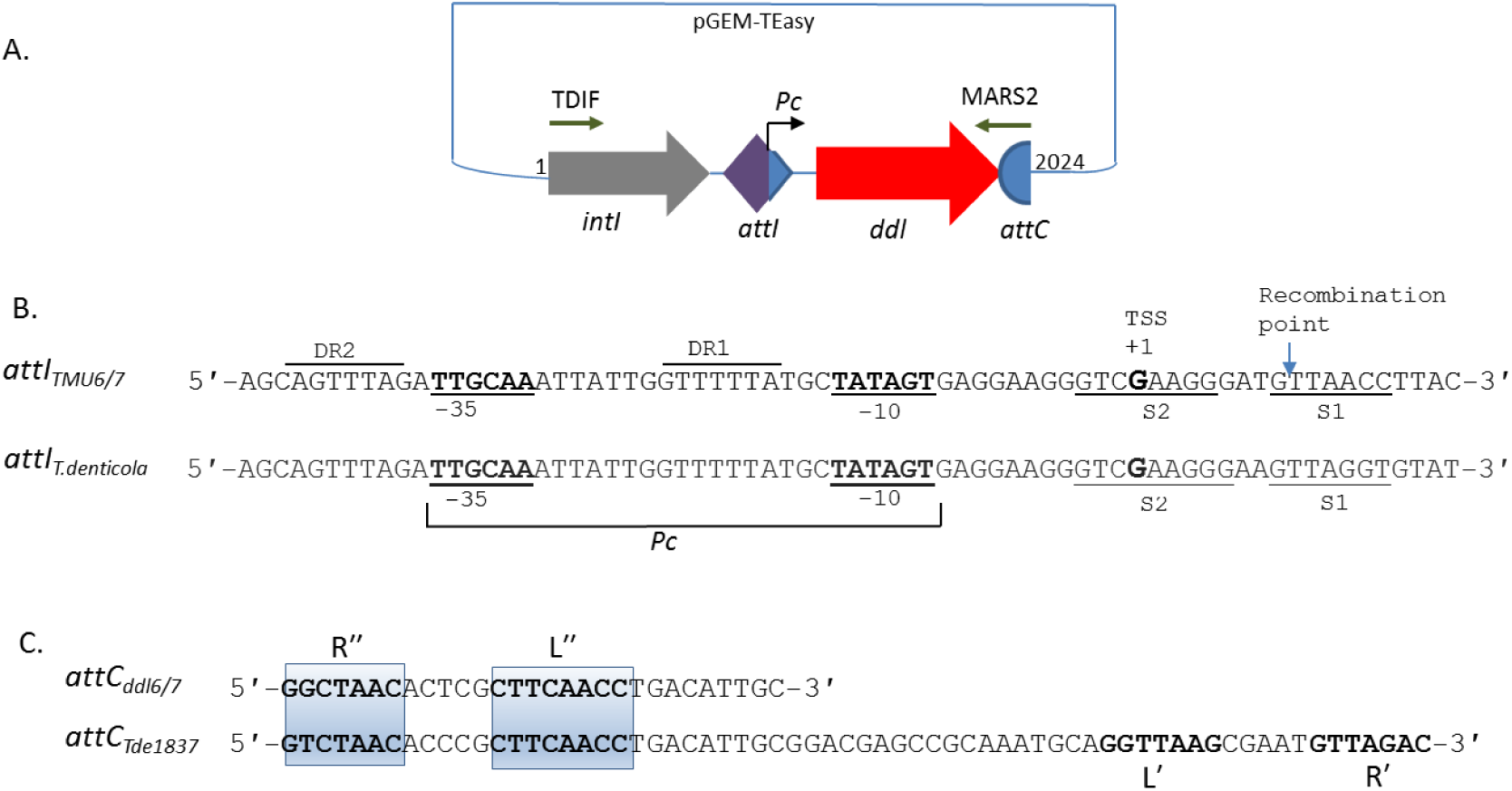
Features of the *ddl* gene cassettes. (A) Genetic organisation of 2,024 bp inserts in the pGEM-T Easy vector; (B) Comparison of putative *attI* sequence preceding *ddl* in the inserts with the putative *attI* of integron of *T. denticola* ATCC 35405 (10). The putative integrase binding sites S1 and S2 as well as DR1 and DR2 are also shown. The putative recombination point G↓TT and the putative transcription start site (TSS) located at the 3′-end of attI are also shown. (C) Comparison of partial sequence of *attC* detected at the 3′-end of the 2,024 bp insert with a typical complete *attC*-associated with Tde1837 of *T. denticola* integron (10).

Two variants of *ddl* (named as *ddl6* and *ddl7*) different from each other by two SNPs at c.490 and c.777 positions of the coding sequences of the genes. The **<u>C</u>**TT (Leu164) and TG**<u>G</u>** (Trp259) codons of *ddl6* were substituted with **<u>T</u>**TT (Phe164) and TG**<u>T</u>** (Cys259) in *ddl7*, respectively.

### The closest homologue of *ddl6* and *ddl7* is a *ddl* located within the accessory genome of *T*.*pedis* B683

The closest homologue of the cassette-located *ddl*s having 98% nucleotide sequence identity (97% identity at the amino acid level) was located on a 5699 bp contig of *T. pedis* B 683 genome (GenBank accession: NZ_AOTN01000179) containing a total of six ORFs. As there was no integrase gene, the association of this contig with an integron, although likely, could not be confirmed. The next closest homologue was Ddl of *Syntrophobotulus glycolicus* (ADY55260) with 55% amino acid identity, followed by *Clostridium sp*. (54%; WP_033164556), *Lachnoclostridium phytofermentans* (53%, WP_029502590) and *Paenibacillus pini* (52%; WP_036650467). Ddl proteins with altered specificity that confer resistance to vancomycin such as VanA (AAA65956), VanB (YP_009076352) and VanC (P29753) also shared low sequence identity (27 to 30%) with Ddl6 and Ddl7. *T. pedis* B683 carries two *ddl* genes, one of which (GenBank accession: AOTN01000124: 8689-9912bp) is most likely the house-keeping *ddl* as it is found in other strains of *T. pedis* including TA4 and TM1 and its G+C content (38.0%) is similar to the genome of *T. pedis* B683 (36.9%). In contrast, the G+C content of the GC located *ddl* is 30.6% suggesting that this second, cassette located *ddl* may have been acquired by horizontal gene transfer by *T. pedis* B683.

### Comparative genomics shows that a likely horizontal gene transfer (HGT) occurred between the *T. pedis* B683 and the host of *ddl6* and *ddl7*

The following three observations suggested HGT between *T. pedis* B683 and the host of *ddl6/ddl7*: i) The identical 40 bp sequences upstream of *ddl6/ddl7* and their homologue *ddl*(GC); ii) the nearly identical (90%) 29 bp downstream sequences containing R″ and L″ sites of *attC* (Fig. 2); iii) the phylogenetic tree constructed using the *ddl*s of different species of *Treponema* showed that the integron-located *ddl6* and *ddl7* and *ddl*(GC) formed a separate clade in the tree (Fig. 3).

**Figure 2.**
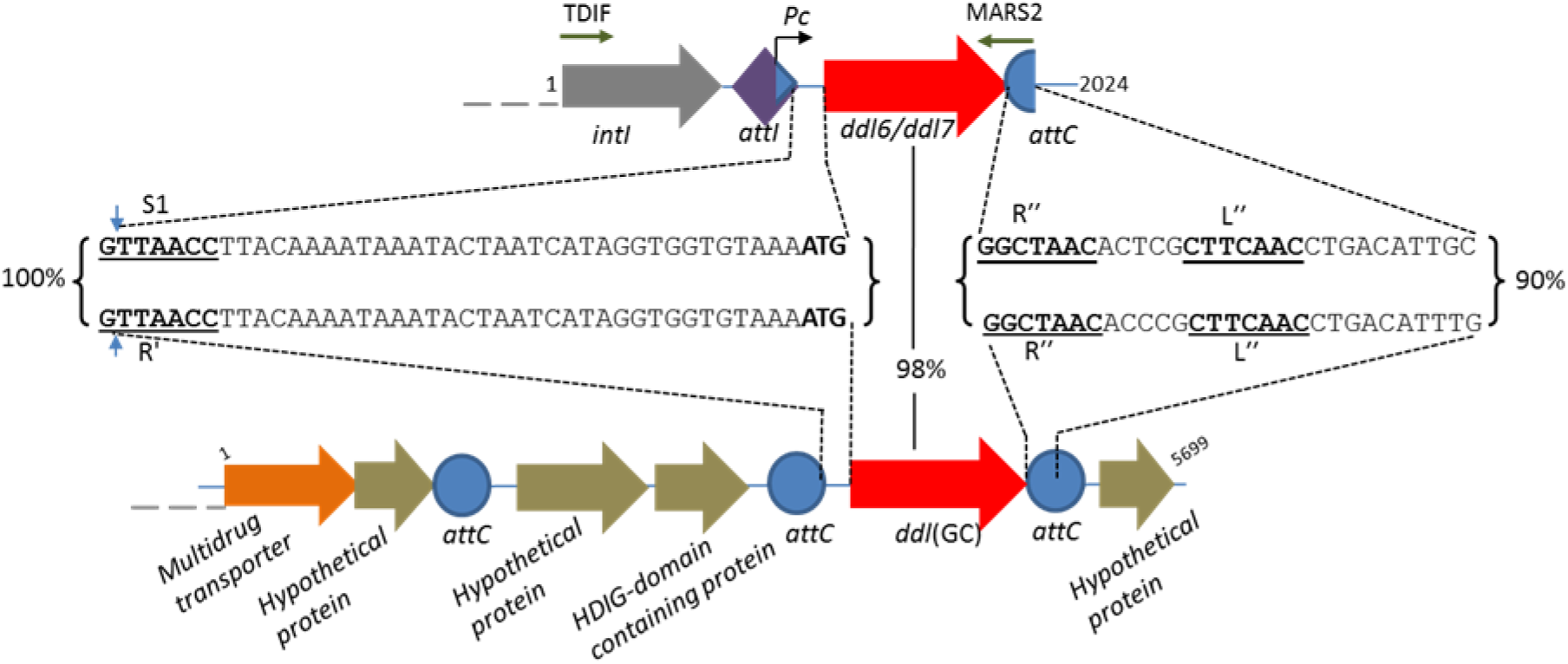
The percentage identity of the flanking sequence of *ddl6/ddl7* (40 bp upstream and 29bp downstream) with their closest homologue, *ddl(GC)* located on a 5699 bp contig of T. pedis B683 genome (GenBank accession: NZ_AOTN01000179). The putative core sites of the *attC* (R″ and L’), the simple integrase binding site (S1) are shown. The recombination points located on S1 and core site, R′ of the *attC* located upstream of *ddl(GC)* (*attC*_*HDIG*_) are marked with the blue arrows. The binding sites for the primers, TDIF and MASRS2, used for amplification of the integrons and associated gene cassettes are also shown. The start codons (ATG) as well as the S1, R′, R″, L″ sites are bolded.

**Figure 3.**
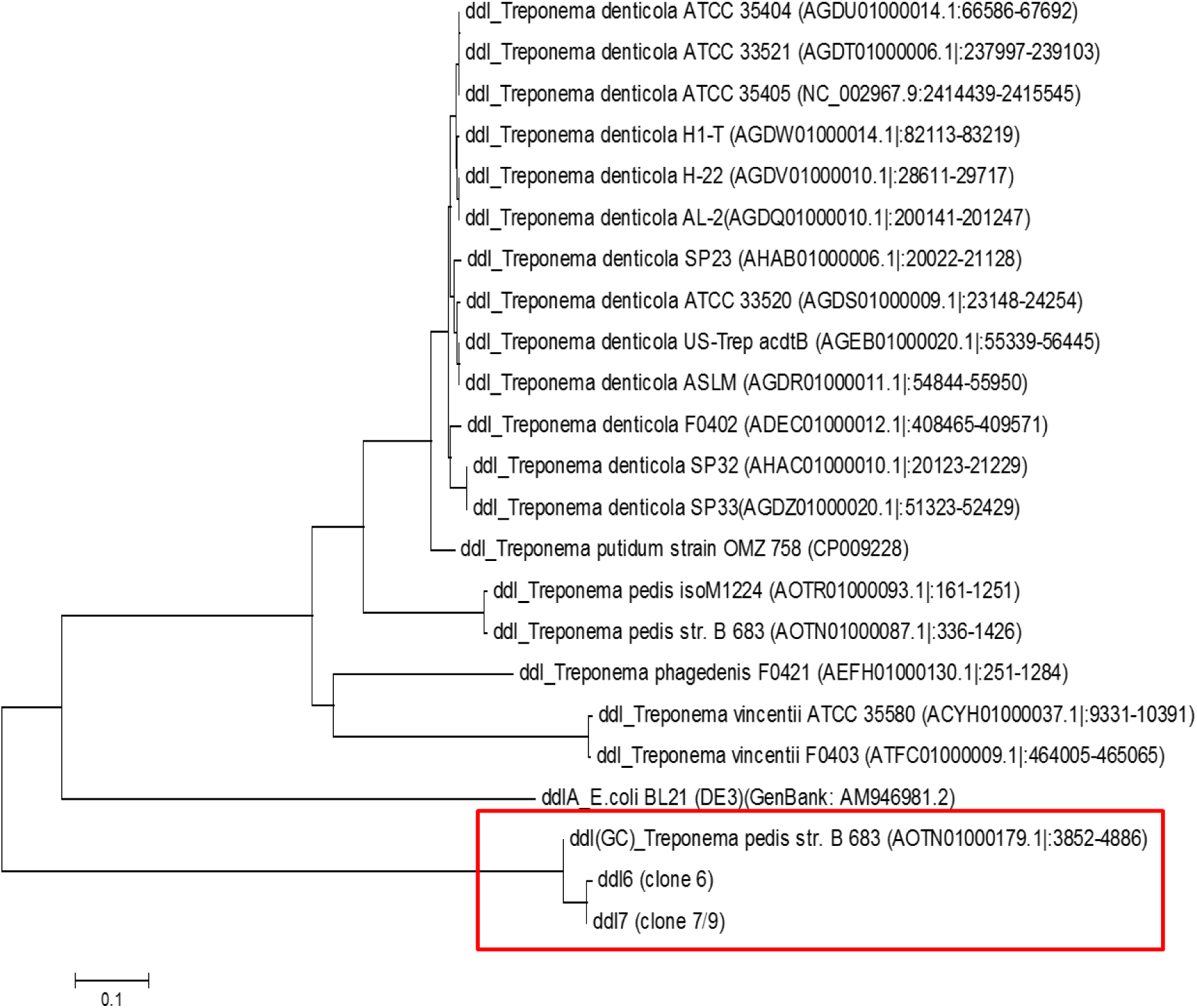
Phylogenetic tree of *ddl6, ddl7* and *ddl(GC)* along with the *ddl*s of different strains of *T. denticola* and other *Treponema* species. The *ddlA* of *E. coli* was used as an outgroup. The evolutionary relationship was inferred using the Neighbour-Joining method. Evolutionary analyses were conducted in MEGA6 (55). The *ddl*s found within gene cassettes are shown in the rectangular box

### Sequence of the upstream region of integron carrying *ddl7* suggests that the putative host of the integron carrying *ddl6* and *ddl7* is a strain of *T. denticola*

To confirm if the initially recovered 2024 bp PCR amplicons were obtained from the *T. denticola* genome, sequence of the *intI* that was further upstream was amplified by PCR. Analysis of a 4,421 bp PCR product showed that it contained a partial ORF encoding a hypothetical protein (1-1,362 bp), a complete ORF encoding IntI (1,266 bp: 1,925 - 3,191 bp), a putative *attI*, the *Pc* promoter followed by a cassette carrying *ddl7* (3,389 - 4,421 bp) (Fig. 4). The complete *intI* had 96.0% identity with the *intI* of *T. denticola* H-22 (GenBank accession no: AGDV01000005:19792-21072). At the amino acid level, the identity ranges between 81-97% with IntI of different *Treponema* sp. Maximum identity was found with the IntI of *T. denticola* AL-2 (97%, EMB42956).

**Figure 4.**
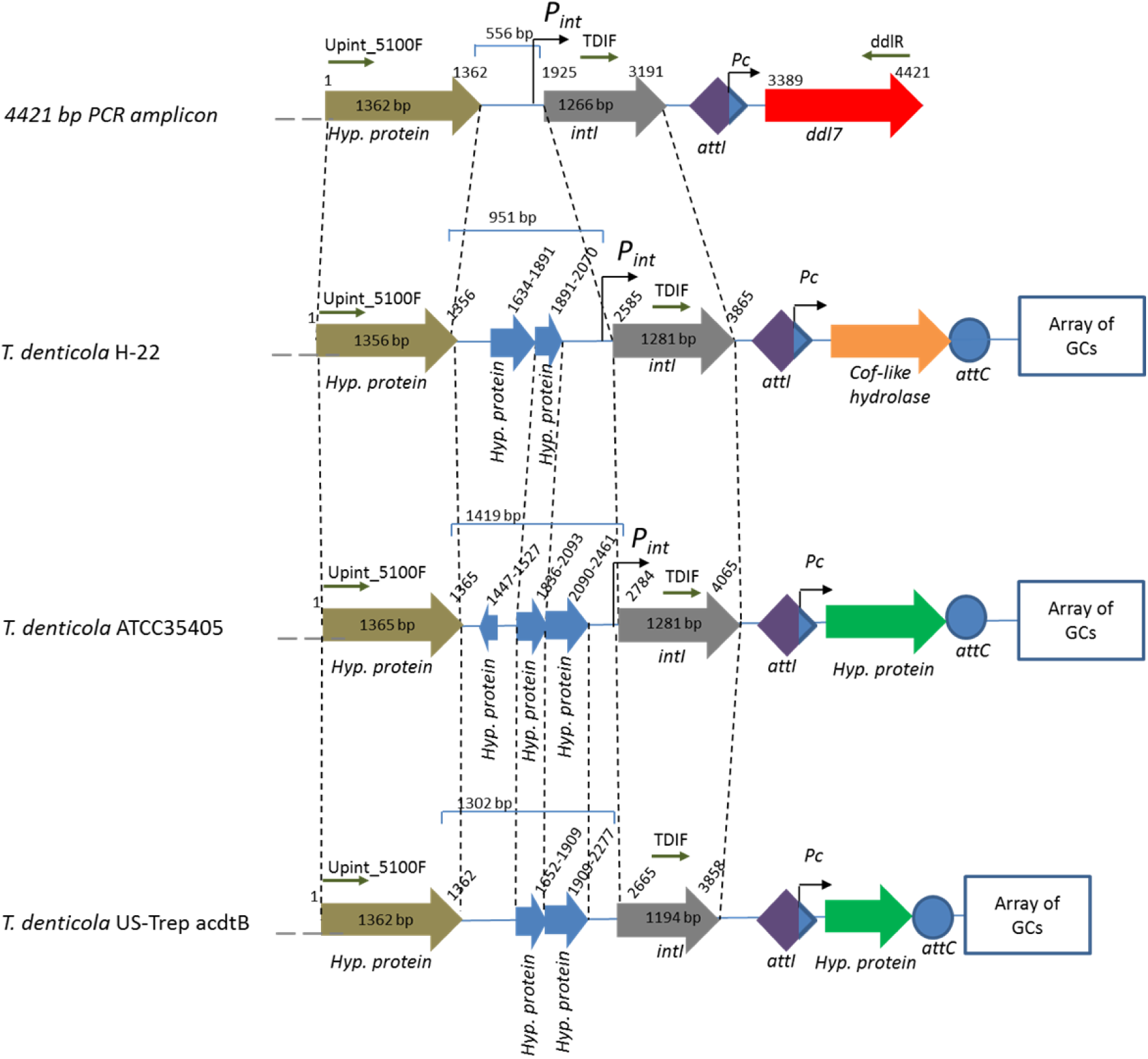
The genetic arrangement of 4421 bp pGEM-T Easy insert carrying *ddl7*, full length *intI* and another partial ORF encoding a hypothetical protein. The primers used to amplify this product are shown with small arrows. The genetic arrangements of the closest homologues including *T. denticola* H-22 (GenBank accession: AGDV01000005:c23656-19631), *T. denticola* ATCC35405 (Genbank accession: AE017226:1873126-1877351) and *T. denticola* US-Trep acdtB (GenBank accession: AGEB01000014.1:3405-5167) have also been shown. The most variable region among the strains was found to be the sequence between the ORF for hypothetical protein and intI.

The genetic organisation of the 4,421 bp amplicon was different from the closely related hits in the NCBI databases (Fig. 4). The non-protein-coding 556 bp sequence located between the partial ORF and the *intI* harbors the putative promoter for the *intI* (*Pi*_*nt*_) with a sequence of 5’-ATGAAT |19 bp| TAAACT-3’. The upstream sequence of the *intI* was analyzed *in silico* for potential LexA binding sites similar to that found in the upstream of *intI* of mobile and chromosomal integrons (13). However, no LexA binding motif could be detected. Additionally, this region contains several inverted repeat and direct repeat sequences as well as a putative transcription terminator sequence.

A phylogenetic tree was constructed to compare the evolutionary relationship of *intI*s associated with the integron carrying *ddl* with the *intI*s of different species of *Treponema*. The tree showed that the *intI* associated with *ddl7* is closely related with other *intI*s of *T. denticola* and distantly related to *T. pedis* (Fig. S1). The results obtained from the *intI* tree as well as the identity of the upstream sequence *intI* with the other strain of *T. denticola* suggest that the likely host of the integron associated with *ddl7* is a species of the genus *Treponema* and most likely a strain of *T. denticola*.

### Expression of *ddl6* and *ddl7* confers resistance to D-cycloserine only, not to vancomycin and beta-lactams

As the overexpression of *ddl* of *M. tuberculosis* and *M. smegmatis* was found to confer D-cycloserine resistance (14), we hypothesised that the expression *ddl6* or *ddl7* in an integron array will result in D-cycloserine resistance, especially as they are detected in the first position due to the efficient expression by the *Pc*. When the minimum inhibitory concentrations (MIC) of D-cycloserine against the surrogate *E. coli* hosts EC126 and EC127 carrying pGEM-T Easy::*intI-attI-ddl6* and pGEM-T Easy::*intI-attI-ddl7*, respectively were tested, a two to four-fold increase of MIC of D-cycloserine was observed (Table 1). As the upstream region of *ddl6* and *ddl7* cassettes carrying the *attI* and *Pc* were 100% identical, we concluded that the fourfold change in the MIC was due to either the SNP c.490 or c.777 or both.

**Table 1.**
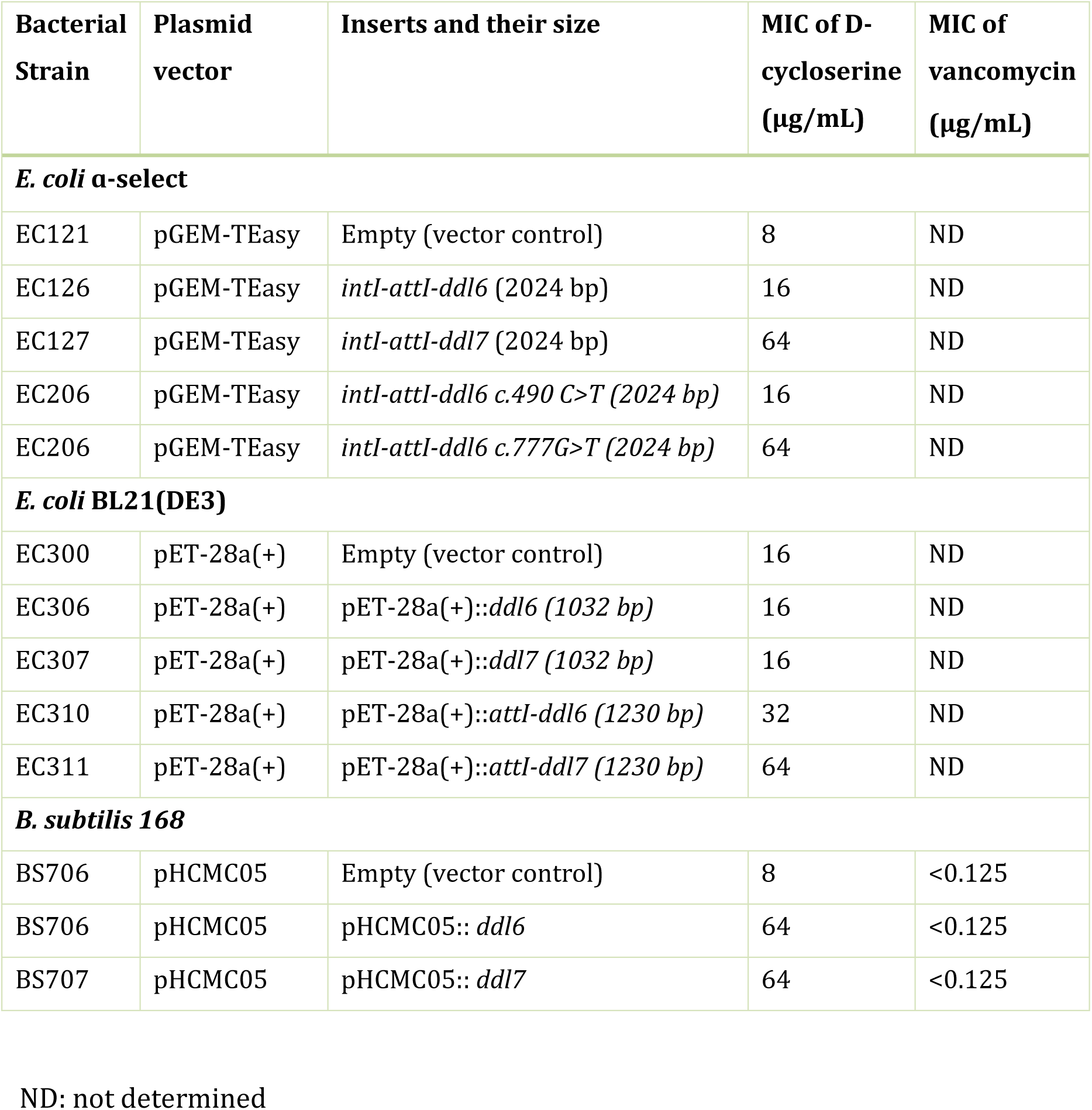
MIC of D-cycloserine against different strains of *E. coli* and *B. subtilis* expressing *ddl*.

To determine if the overexpression of *ddl* in gram-positive bacteria can also confer resistance to D-cycloserine, vancomycin and β-lactam antibiotics, the 1032 bp coding sequence of *ddl6* and *ddl7* was cloned into a medium-copy expression vector, pHCMC05 and transformed into *B. subtilis* 168. The MIC of D-cycloserine against the strains BS706 and BS707 which carry pHCMC05::*ddl6* and pHCMC05::*ddl7* vectors, respectively was found to increase by eight-fold (64 μg/mL) compared to the BS700 strain that carries the empty plasmid (8 μg/mL) (Table 1). However, no difference in the MIC of D-cycloserine was observed between BS706 and BS707 carrying *ddl6* and *ddl7*, respectively, when the expression of the genes was induced by IPTG. No resistance to vancomycin was observed (Table 1). Nor was there any resistance to penicillin G, amoxicillin, oxacillin and ampicillin against both *E. coli* and *B. subtilis* strains. This suggests that overproduction of Ddl does not confer cross-resistance to antibiotics targeting cell-wall biosynthesis.

### Site-directed mutagenesis confirmed that the SNP at c.777 was responsible for the alteration of MIC of D-cycloserine

To identify which SNP of *ddl6* and *ddl7* was responsible for altering the susceptibility of *E. coli* to D-cycloserine, the nucleotides at position c.490 and c.777 of *ddl6* were changed to the corresponding nucleotides of *ddl7* by site-directed mutagenesis. MICs of D-cycloserine against *E. coli* carrying the constructs were determined and no change in the susceptibility was observed by the c.490 C>T substitution in *ddl6*, however, the G>T substitution which alters Trp259 to Cys259 (W259C) increased the MIC fourfold which is similar the MIC of the strain carrying *ddl7* (Table 1).

### The integron-encoded Ddls are functional

The biological activity of Ddl6 and Ddl7 was confirmed using purified proteins (Fig. 5A). The release of the inorganic phosphate (Pi) in the Ddl-catalyzed reaction was assayed and both variants of Ddl catalyse the release of a significant amount of Pi into the reaction compared to the control (P>0.05) (Fig. 5B). This indicates that ATP is consumed in the reaction and Pi is released. Furthermore, paper chromatography analysis of the Ddl-catalysed reaction product showed that both proteins catalyze the ligation of D-ala resulting in the synthesis of D-ala-D-ala dipeptide (Fig. 5C). The substrate specificity assay indicated that they could only catalyse the synthesis of D-ala-D-ala dipeptide, but not D-ala-D-ser or D-ala-D-lac.

**Figure 5.**
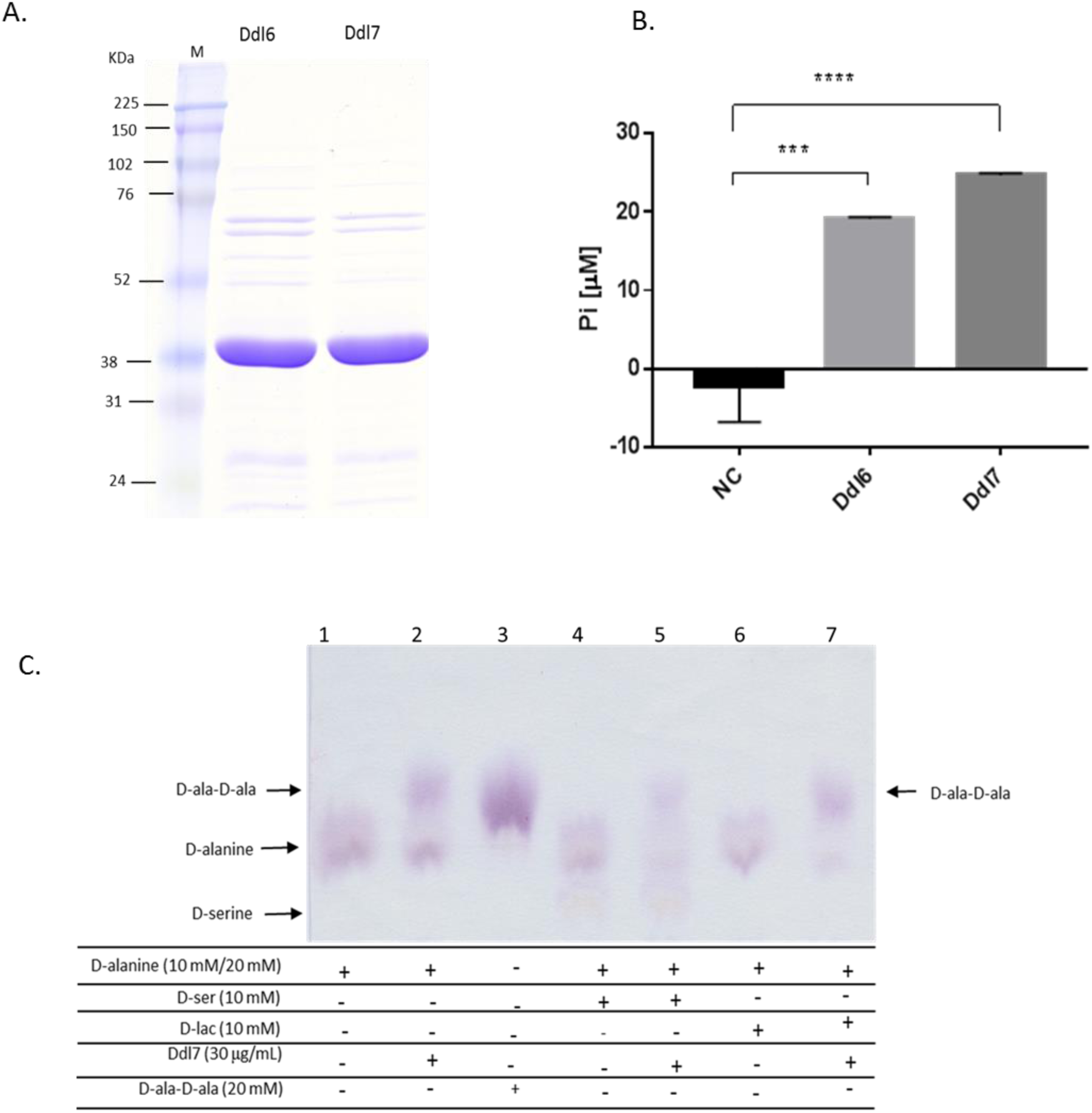
Purification of Ddl and determination of the functional activity Ddl. (A)SDS-PAGE of purified Ddl6 and Ddl7. (B) Release of inorganic phosphate in the reactions catalysed by Ddl6 and Ddl7 in the presence of 20 mM D-alanine. The differences of the release of Pi in between the negative controls (no Ddl) and Ddl catalysed reactions were analysed by one-way ANOVA (***P<0.001; ***P<0.0001). (C) Ascending paper chromatography to detect the formation of D-ala-D-ala and D-ala-D-ser dipeptide as well as D-ala-D-lac depsipeptide. - In the first two reactions (lane 1 and lane 2) 20 mM D-alanine was used, whereas in other reactions it was reduced to 10 mM and supplemented with 10 mM D-serine or D-lactate.

### The predicted 3D structures of Ddl6 and Ddl7 provided information on alteration of resistance phenotype and substrate specificity

To determine if the alteration of Trp259 of Ddl6 to Cys259 of Ddl7 caused a conformational change, 3D models of both Ddl6 and Ddl7 were constructed using I-TASSER (15) (Fig. S2). When the structures were superimposed using TM-align (16), they superimpose near perfectly with a TM score of 0.97. However, a notable predicted change in the conformation of the omega-loop region was observed likely resulting from the substitutions at 164 and 259 positions of Ddl6 and Ddl7 (Fig. S3).

Three putative domains of Ddl6 and Ddl7 were identified by comparing predicted 3D structures of Ddl6 and Ddl7 with the crystal structure of D-ala-D-ser ligase (VanG, PDB code: 4FU0) (17), DdlB of *E. coli* (18), VanA of *E. faecium* (19) and Ddl of *S. aureus* (20). The predicted N-terminal domain of Ddl6 and Ddl7 runs from the N-terminus to Gly125, central domain from Ser126 to Gly219 and C-terminal domain from Arg220-Ile340. The putative omega-loop, which plays an important role in catalytic activity and substrate specificity of Ddl proteins, was located on the C-terminal domain of Ddl6 and Ddl7 and runs from Lys240-Ser255. The Trp259 of Ddl6 whose substitution with Cys259 causes a fourfold increase in D-cycloserine resistance was found to be located at the end of the 4^th^ β-sheet (residues Asn256-Ile258) of the C-terminal domain and outside of the omega-loop (Fig S2). The amino acid residues (Ser150 and Tyr216) of *E. coli* DdlB which are conserved in the Ddls that can only produce D-ala-D-ala residues were also conserved in Ddl6 and Ddl7 with the corresponding residues of Ser184 and Tyr250 (Fig. S4).This supports the results of the *in-vitro* assays and explains why it could produce only the D-ala-D-ala dipeptide, not D-ala-D-ser or D-ala-D-lac.

Although the integron-encoded Ddls were not found to confer vancomycin resistance and did not carry the amino acid residues for altered substrate specificity (D-Ser or D-Lac), among the closest homologues in PDB that matched with the predicted 3D structures of Ddl6 and Ddl7, four were related to vancomycin resistance including VanG (PDB code: 4FU0) and VanA (PDB code: 1E4E) with high TM scores (Table S1).

### Molecular Dynamics simulations post-analysis provides insights into the mechanism of alteration of MIC of D-cycloserine due to W259C mutation in Ddl6

The results of 50 ns molecular dynamics (MD) simulations of wild-type Ddl6 and its W259C derivative in complex with D-cycloserine (Fig. 6) showed that the ligand is released from the recombinant protein after approximately 20 ns, however, the complex with the wild-type protein remains stable in the binding site during the course of the MD simulations.

**Figure 6.**
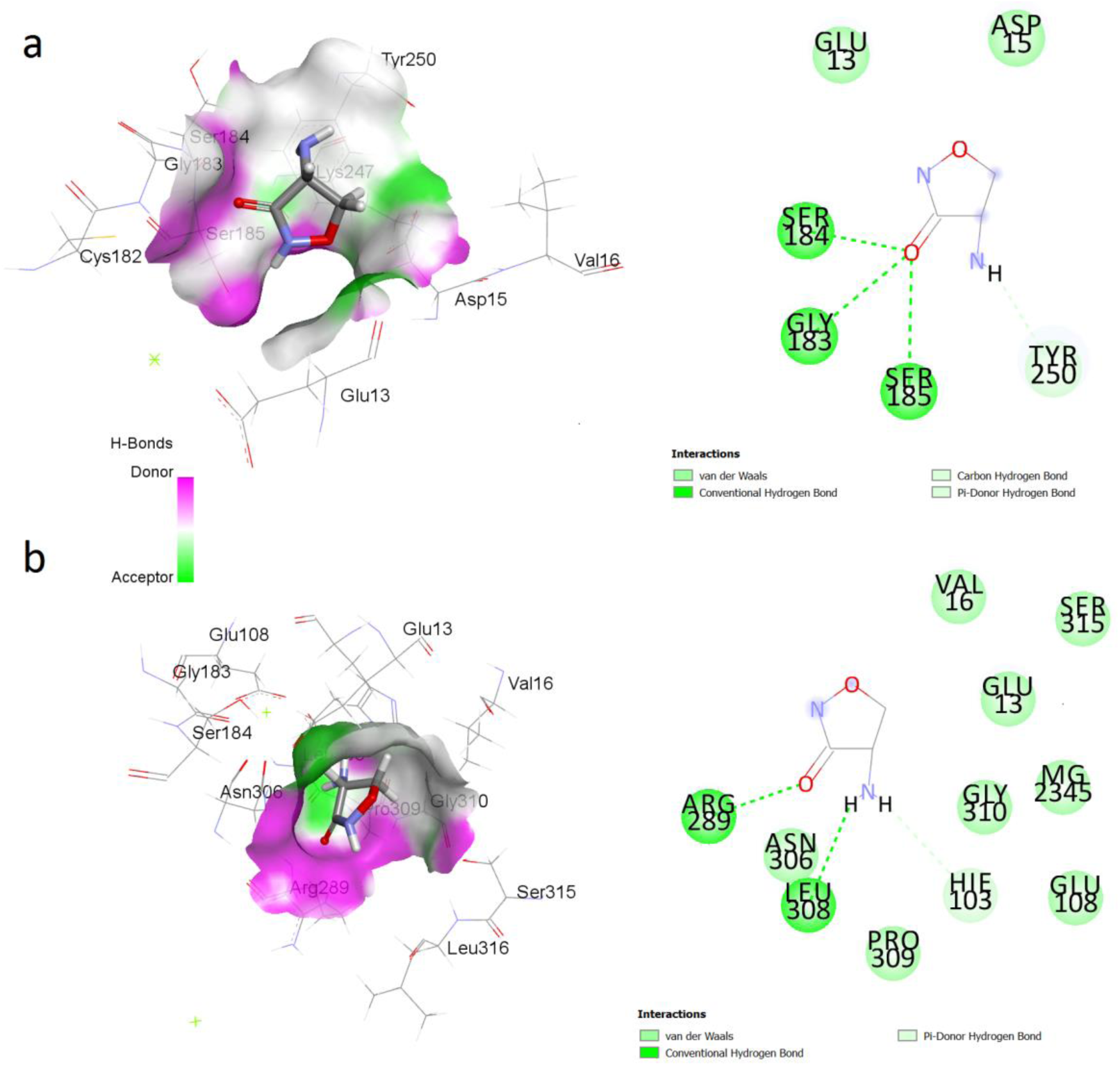
2D (right panels) and 3D (left panels) structures of the binding sites of the W259C mutant of Ddl6 (a) and wild type Ddl6 (b) in interaction with D-cycloserine in average structures extracted from MD trajectories. After placing in the same location at the beginning of the MD simulations, both ligands were found in the adjacent grooves in the same binding site of their corresponding average structures.

The movement of the ligand from the protein was studied by monitoring the distances (Fig S4) between the centre of D-cycloserine and Trp259 of the wild-type Ddl6 and Cys259 of the Ddl6 W259C mutant during the course of the simulation.

The distance between the centre of D-cycloserine and Cys259 of the mutant protein progressively increased suggesting a movement away from the binding site, whereas the distance between the centre of D-cycloserine and Trp259 of the wild-type protein remained relatively stable throughout the simulation (Fig. S5)

The relative binding free energy values calculated from the MD simulations trajectories further supported this observation as the complex of wild-type Ddl6 and D-cycloserine was more favorable (−14.47 kcal/mol) than the complex of D-cycloserine and Ddl6 W259C mutant complex (−12.15 kcal/mol) (Table S2). Videos of the MD simulations showing the movement of the ligands are available in Video S1 and Video S2.

### Kinetic analysis of pure proteins showed that Ddl7 is more active than Ddl6

The integron-encoded Ddl6 and Ddl7 were characterised kinetically and the kinetic parameters were compared with DdlTd (the Ddl encoded by house-keeping *ddl* of *T. denticola*) DdlAEc as positive control. The K_m,D-ala2_ of Ddl6 and Ddl7 was determined as 0.39 and 0.54mM, respectively (Table 2), thus, the values of K_m,D-ala2_ could not explain the alteration of D-cycloserine phenotype between these two variants. However, the turnover number (K_cat_) of Ddl7 (3.85 s^-1^) was approximately three-fold higher than Ddl6 (1.20 s^-1^), which nearly matches the fourfold increase of MIC of D-cycloserine for Ddl7 as compared to Ddl6.

**Table 2.** Steady-state kinetic parameters of Ddl6, Ddl7, DdlTd and DdlEc.

| **Proteins** | **K<sub>m</sub>, ATP (μM)** | **K<sub>cat</sub> (s<sup>-1</sup>)** | **K<sub>m</sub>, D-ala2 (mM)** | **K<sub>cat</sub>/K<sub>m</sub>, D-ala2 (M<sup>-1</sup> s<sup>-1</sup>)** |
| --- | --- | --- | --- | --- |
| Ddl6 | 54.96 | 1.19 | 0.39 | 3008.85 |
| Ddl7 | 43.28 | 3.85 | 0.55 | 7010.32 |
| DdlTd | 19.08 | 3.74 | 0.45 | 8294.55 |
| DdlAEc | 48.99 | 6.43 | 0.52 | 12517.21 |
| DdlAEc* | 38* | NR | 0.55* | NR |

\* The values of DdlAEc reported by Zawadzke, Bugg and Walsh (32). NR, not reported

### Molecular docking analysis showed that flavonoids could be responsible for selection of the *ddl6/ddl7* within the first position of the integron

We also tried to identify the likely selection pressure that could drive the evolution of *ddl*s within integrons and maintaining them as the first GC. As the use of D-cycloserine is restricted only for the MDR and XDR *Mycobacterium tuberculosis*, it is very unlikely to be responsible for this. We assumed that this selective pressure is common for both countries from which *ddl* containing strains were isolated (Bangladesh and the UK), as the *ddls* were ubiquitous within samples from both geographic locations. Candidates for providing this selective pressure are two plant flavonoids, quercetin and apigenin, which are abundantly present in many teas, fruits and vegetables and inhibit Ddl (21). Molecular docking was used to see if these flavonoids can bind to Ddl within the predicted D-alanine and/or ATP binding sites of Ddl6 and Ddl7. Salvicine (a terpenoid isolated from a Chinese medicinal plant, *Salvia prionitis*) was used as a positive control for the docking because this terpenoid was identified as a potent inhibitor of Ddl of *T. pallidum* using molecular docking (22). GOLD molecular docking showed that quercetin and apigenin can dock to both D-alanine (Fig. 7; Table S3) and ATP binding sites (Fig. S6; Table S3) of Ddl6 and its mutants with thus, the integron-encoded Ddls may play a role in both the sequestration of these inhibitors, lowering the intracellular concentration and allowing the house-keeping Ddl to perform its physiological functions and also take on the role of the house-keeping Ddl during cell wall development.

**Figure 7.**
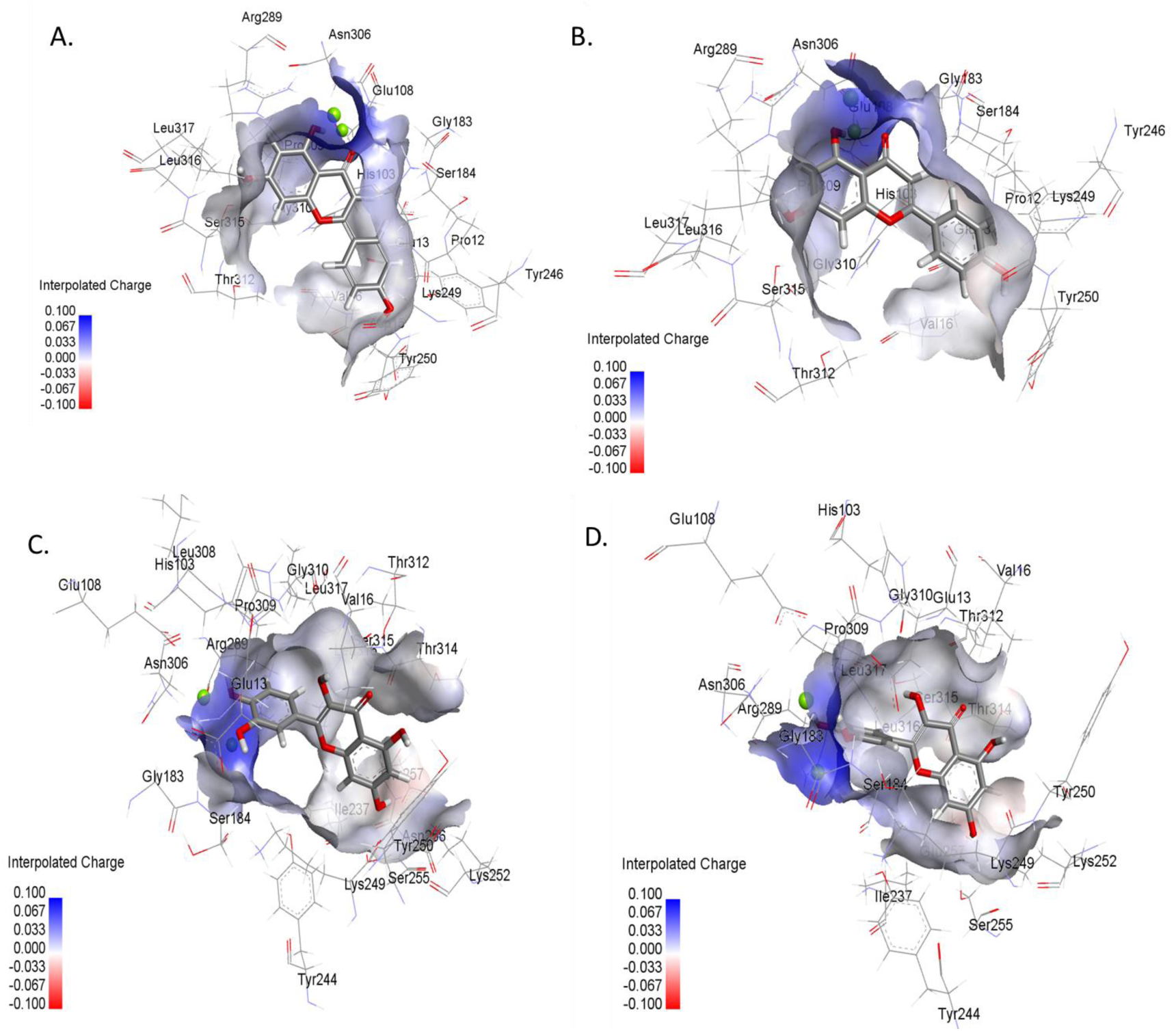
Mode of interaction of apigenin and quercetin with the D-alanine binding sites of wild type Ddl6 and its W259C mutant. A) Apigenin with WT Ddl6, B) Apigenin with W259C mutant, D) Quercetin with WT Ddl6 and D) Quercetin with W259C mutant.

## Discussion

This study reports for the first time the discovery of integron GCs encoding two novel isoforms of Ddl (Ddl6 and Ddl7) derived from the oral cavity microbiota of healthy humans in the UK and Bangladesh. The ORFs encoding *ddl* were located immediately downstream of *intI* and oriented at the same direction as on reverse integrons. All of the genetic characteristics of an integron and its associated gene cassettes were found within the amplicon, including the presence of a putative *Pc*, an *attI* region harbouring putative integrase binding sites (S1 and S2), a putative *attI/attC* recombination site, a putative RBS located 6 bp upstream of the start codon of the *ddl*s as well *attC* core sites (R″ and L″)(5, 8, 23, 24).

The closest homologue of *ddl6* and *ddl7* with approximately 98.0% nucleotide identity was located on a whole genome sequence (WGS) contig of T. *pedis* B683 and is one of two copies of *ddl* within this genome and is not present in the genome of other *T. pedis* strains, thus it is likely part of the accessory genome. *T. pedis* is an animal pathogen that causes digital dermatitis in cattle and necrotic ulcers in pigs (25). B683 (GenBank (NZ_AOTN00000000.1) was isolated from a shoulder ulcer of pigs. The evidence showed that there is a likely genetic exchange between *T. pedis* and the host of *ddl6/ddl7*, likely to be *T. denticola*, which may involve one or more transfer intermediates.

Analysis of the upstream sequence of the integron carrying *ddl6/ddl7* suggests that the likely host of the integron is a strain of *T. denticola* which carries two copies of *ddl* in its genome: one as an integron gene cassette and another as house-keeping gene (this house-keeping *ddl* of *T. denticola* ATCC35404 showed only 46.0% nucleotide identity to *ddl6 and ddl7*). *T. denticola* is one of the most comprehensively characterised species among the 49 species of *Treponema* in the human oral cavity (26). It is one of the members of the ‘red complex’ associated with clinical progression of periodontitis (27) and was capable of transferring genes to different genera of bacteria in dental plaque (28). It is the only bacterial species in the oral cavity that has been found to carry integrons (10) although we have recently demonstrated other bacterial species are also likely to contain integrons (29). From the *in silico* analysis of the metagenomic datasets of the NIH Human microbiome project, a total of 826 gene cassettes related to *Treponema sp*. were detected in the oral cavity (30). The closely related strains of *T. denticola* shared a high nucleotide identity (>80%) of their integron gene cassettes (31).

Our results provide novel insights into the evolutionary mechanisms that have facilitated the acquisition of a second copy of *ddl* on an integron in the original host of *ddl6* and *ddl7*. Zawadzke, Bugg and Walsh (32) proposed two plausible explanations for the duplication of *ddl* in *E. coli*: a) one of the two genes (*ddlA*) is expressed constitutively whereas the expression of the other gene (*ddlB*) is controlled tightly and coupled with the expression of other cell-wall biosynthesis genes; b) the D-ala-D-ala dipeptide may be used for another pathway other than for peptidoglycan synthesis. Both explanations proposed by Zawadzke, Bugg and Walsh (32) for the duplication of *ddl* in *E. coli* are likely not applicable here because, unlike the duplicated *ddlA and ddlB* genes of *E. coli*, the *ddl* cassettes are likely to have been acquired by HGT and thus, they may be related to gaining a new function (resistance to antimicrobials) in addition to, or rather than, providing increased amount of the gene product (33). The kinetic analysis of the purified proteins (Ddl6, Ddl7 and DdlTd) also indicates that the *ddl*s are interchangeable with respect to their core function of cell wall assembly because, the K_cat_ (s^-1^) of DdlTd was very close to Ddl7 thus, the acquisition of a second copy of *ddl* within integrons provide resistance by titrating out intracellular antimicrobials plus complementation of the original Ddl function.

We found that the ectopic expression of *ddl6* and *ddl7* confers resistance to D-cycloserine. As Ddl is a target of D-cycloserine, overproduction of Ddl is likely to facilitate resistance by mechanisms similar to D-cycloserine resistance in *Mycobacterium* which is mediated by high expression of both Ddl and Alr (14, 34). The other known mechanisms of D-cycloserine resistance in *Mycobacterium* were the non-synonymous mutations in Alr including S22L, K133E, L89R, M319T, Y364D and R373G (35, 36) as well as loss of function mutation in *alr* (36). In *E*.*coli* the mutation in *cycA* (37), *dadA* and *pnp* (38) confer resistance to D-cycloserine. The mutations in a penicillin binding protein (PBP4) homologue in *M. smegmatis* confer resistance to D-cycloserine and vancomycin (39). Although several mutations in *alr* have been implicated in D-cycloserine resistance, so far no mutations in *ddl* were found to be associated with alteration of D-cycloserine resistance by the host. Moreover, several functional metagenomic analyses also reported that ectopic expression of various *ddl* also conferred resistance to D-cycloserine (40-43).

While the expression of both integron encoded *ddl*s conferred resistance to D-cycloserine, the expression of *ddl7* had a four-fold higher MIC compared to *ddl6*. Site-directed mutagenesis confirmed that a single substitution at c.777, which alters Trp259 of Ddl6 to Cys259, was responsible for this. Using MD simulations, we have shown how this mutation is predicted to alter the binding affinity of the ligand D-cycloserine to the active sites of the proteins. The wild-type Ddl6 with a tryptophan residue at its 259-position had more stable binding to D-cycloserine, therefore less D-cycloserine was required to occupy active sites which in turn decreased the MIC. When the tryptophan residue was changed to cysteine (the residue at 259 for Ddl7), binding between the mutant Ddl6 and D-cycloserine complex was comparatively less stable and thus, more antibiotic will be needed to occupy the active sites which causes an increase in the MIC.

The differences in the properties of cysteine and tryptophan could be responsible for changing the binding of D-cycloserine to the active sites of Ddl. Cysteine is a polar amino acid and its reactive thiol (−SH) group is essential for the bioactivity of many proteins (44). Moreover, cysteine residues form disulphide bridges which are crucial determinants for the tertiary protein structure (45). In contrast, tryptophan is an aromatic non-polar hydrophobic amino acid. Although it plays an important role in protein folding, it is not reactive like cysteine. Thus, due to the differing nature of both amino acids, Trp→Cys substitution in Ddl6 is likely to have a major impact on its structure and function by affecting the ligand binding and catalytic activity.

Among the closest structural homologues of Ddl6/Ddl7 in PDB, four were related to vancomycin resistance which include VanA of *E. faecium*, VanG of *E. faecalis* and D-alanine-D-lactate ligase of *L. mesenteroides*. The high TM-scores (>0.9 for VanG and >0.85 for VanA) also indicate close 3D structural similarity. It is known that the function of a protein is more related to the structure rather than the sequence (46). The inclination of the predicted 3D structures of Ddl6 and Ddl7 towards vancomycin resistant determinants suggests that if the corresponding residues of VanG or VanA, which are responsible for binding and positioning D-ser (VanG-type) or D-lac (VanA-type) instead of D-ala2, are substituted to the corresponding residue of Ddl6 and Ddl7 (Tyr250), they could gain D-ala-D-ser dipeptide or D-ala-D-lac depsipeptide ligase activity. As the first few GCs within integrons are mostly more efficiently expressed, the Ddls encoded by integrons that acquire D-ala-D-lac or D-ala-D-ser ligase activity by mutation, may outnumber by the D-ala-D-ala and therefore may not need alternative vancomycin-resistance associated genes such as those found in vancomycin-resistant enterococcus (VRE) (47). The *E. coli* DdlB gained a weak D-ala-D-lac depsipeptide activity following Tyr216 and Ser150 substitutions with phenylalanine and alanine of LmDdl2, respectively (48).This supports our hypothesis that if the corresponding residue of VanA, VanG or LmDdl2 are substituted with the Tyr250 of Ddl6/Ddl7, they could be able to confer resistance to vancomycin.

We also suggest a hypothesis for evolution of *ddl* within an integron cassette within the first position. D-cycloserine is currently only used in the treatment of MDR and XDR-tuberculosis and there is no history of environmental use of D-cycloserine in agriculture. Additionally, none of the volunteers received this drug (or any other antibiotic), at least in the three months prior to saliva collection. Therefore, it is very unlikely that D-cycloserine represents the driving force for this event. We hypothesize that the dietary flavonoids or the inhibitory small molecules present in our diet that can bind Ddl driving the acquisition and selection of *ddl* within integrons. Some flavonoids including apigenin, quercetin and catechin are known to have antibacterial activity (49) and are found in fruits and vegetables as well as widely consumed beverages such as tea. Apigenin and quercetin has been shown to inhibit the activity of Ddl of *H. pylori* (21) by competitively inhibiting the binding of ATP to ATP-binding sites. MD analysis performed in this study supported the findings and demonstrated that these flavonoids are likely to bind to both D-ala and ATP binding sites of Ddl6 and Ddl7. Thus, the excess Ddl produced by the efficient expression under the control of *Pc* could confer a selective advantage on the host titrating out inhibitory flavonoids (Fig. 8). This hypothesis requires testing experimentally which is now under way.

**Figure 8.**
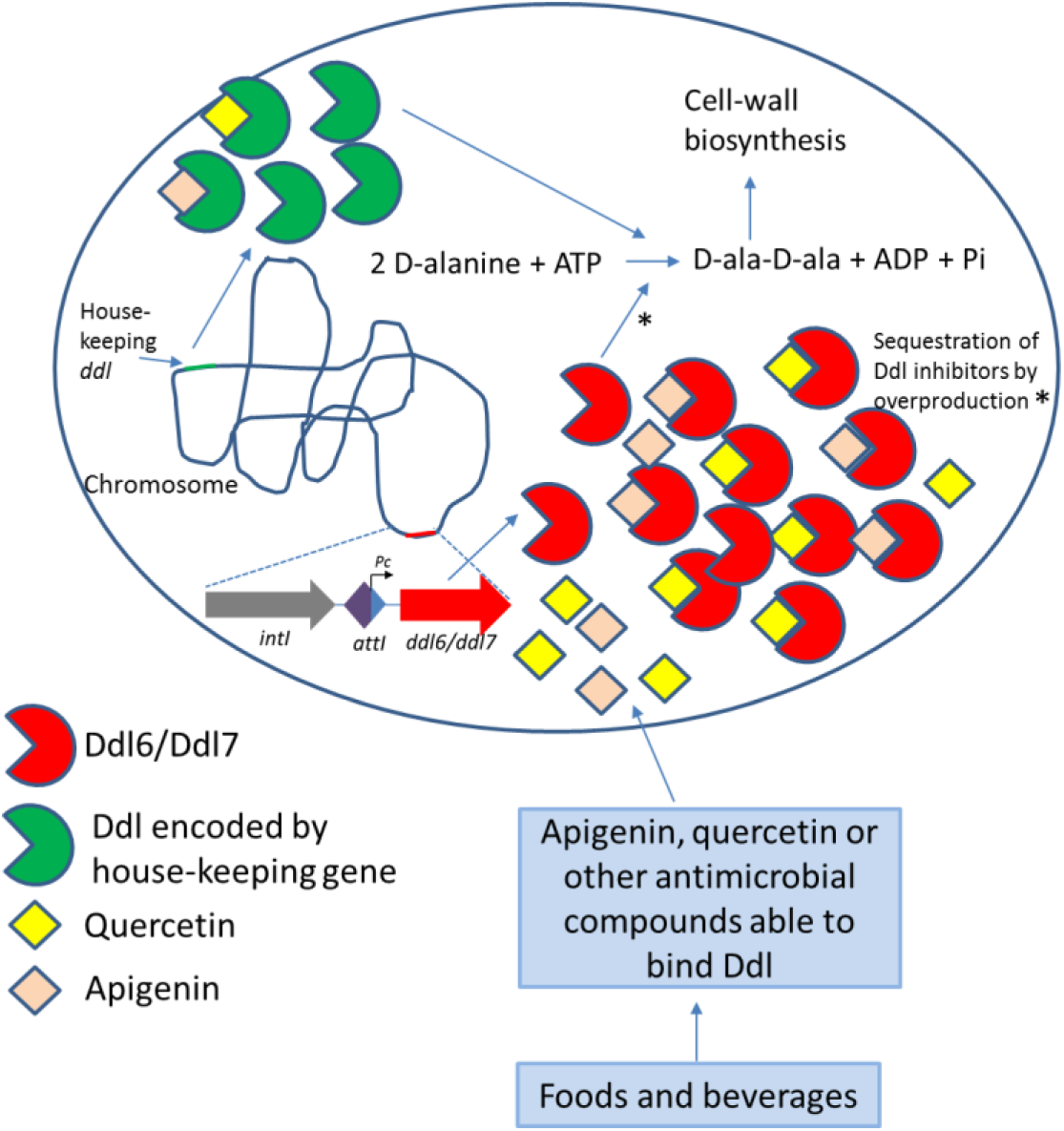
Proposed model for plant flavonoids promoting the selection of *ddl*s and their maintenance within the first GC of the integron of *T. denticola*. The integron encoded Ddl6/Ddl7 could play a dual role (marked with asterisk): sequestration of flavonoids or other molecules that are able to bind Ddl and catalysing the formation of D-ala-D-ala dipeptide in presence of inhibited Ddl encoded by the house-keeping genes.

The results show that *ddl*, which is generally considered a house-keeping gene has moved onto an integron and can be found in the first position, with the most likely selection pressure being dietary compounds with antimicrobial activity. Because these genes also confer resistance to D-cycloserine and are potentially one mutation away from vancomycin resistance, the study also sheds light on potential environmental drivers of clinically relevant antimicrobial resistance.

## Materials and Methods

### Source of chemicals, reagents and antibiotics

Chemicals, solvents, culture media and antibiotics were purchased from Sigma-Aldrich Ltd (Dorset, UK), BDH (UK) and Life Technologies (USA) unless stated otherwise. Plasmid preparation kits were obtained from Qiagen (UK). Restriction enzymes were obtained either from New England Biolabs (UK) Ltd or from Promega (UK). The EnzChek Phosphate assay kit and Pierce BCA Protein assay kits were purchased from Life Technologies (USA). *E. coli* α-Select competent cells were obtained from Bioline Reagents Ltd (UK). The primers were obtained from Sigma-Aldrich (UK).

### Bacterial strains and plasmids

The pGEM-TEasy vector (Promega) was used for constructing the metagenomic library of PCR amplicons of integrons and gene cassettes. *E. coli* α-select silver efficiency (Bioline) was used to manipulate the pGEM-TEasy library of gene cassettes. pET28a (Novagen) and *E. coli* BL21 (DE3) (New England Biolabs) were used for cloning and expression of *ddl*s and purification of corresponding His-tagged proteins, respectively. *Bacillus subtilis* 168 and pHCMC05 (Bacillus Genetic Stock Centre) were used for cloning and expression of *ddl6* and *ddl7* to determine the susceptibility of D-cycloserine and vancomycin in Gram-positive bacteria. *E. coli* strains were cultured in LB medium at 37°C. *Bacillus subtilis* 168 was cultured in Brain-heart infusion (BHI) medium unless stated otherwise. The bacterial strains were preserved at -80°C in 30% (v/v) sterile glycerol.

### Detection of *ddl* gene cassettes in the metagenomic DNA obtained from saliva

The collection of saliva samples, preparation of metagenomic DNA from pooled saliva from UK and Bangladesh and detection of integrons and gene cassettes in the library PCR amplicons have been described previously (29). Briefly, the PCR products produced using the forward integrase primer (TDIF) and reverse *attC*-based primer (MARS-2) were cloned into pGEM-TEasy and transformed into *E. coli* alpha select cells. Transformants were randomly picked, plasmids were prepared from 5 ml overnight culture using Miniprep kits (Qiagen) and the plasmids were sequenced using M13F-M13R primers (Table S4).

### Determination of the upstream region of integron carrying *ddl6* and *ddl7*

The forward primer, Upint_5100F (Table S4) was coupled with the reverse primer, ddlR to amplify the full length of *intI* and its upstream region. Q5® High-Fidelity 2X Master Mix (NEB, UK) was used for PCR using the pooled metagenomic DNA of UK saliva samples as template. The expected size PCR products were purified by gel extraction using QIAquick Gel Extraction Kit (Qiagen, Germany) and A-tailed following the protocol for cloning blunt-ended PCR products described in pGEM®-T Easy Vector Systems Technical Manual (Promega, UK). An aliquot of A-tailed PCR products was cloned into pGEM-T Easy and then transformed into *E. coli* alpha select cells. The transformants were selected on LB agar plates supplemented with ampicillin, IPTG and X-gal. Plasmids were prepared and the inserts were sequenced.

### Site-directed mutagenesis

The Phusion Site-Directed Mutagenesis Kit (Thermo Scientific, USA) was used to construct two different mutants of *ddl6: ddl6* c.490 C>T and *ddl6* c.777 G>T using the pGEM-T Easy::*intI-attI-ddl6* plasmid as template. The phosphorylated mutagenic primers used for constructing the mutants are listed in the Table S4. After confirming the success of the mutagenesis reactions, the mutant plasmids were transformed to *E. coli* α-select cells and MIC of the transformants were determined using the agar dilution method.

### Sub-cloning of *ddl6* and *ddl7* into pHCMC05 and transformation into *B. subtilis*

The genes for Ddl6 and Ddl7 were amplified by PCR using the pGEM-TEasy vectors with the 2024-bp insert containing *ddl6* and *ddl7* as template (see Table S4 for primers used). The genes were cloned into *BamH*I and *Xba*I sites of pHCMC05 (*E*.*coli-B. subtilis* shuttle vector) using the primers TddlF and TddlR (Table S4). Chemically competent *B. subtilis* 168 cells were prepared and transformed with the recombinant pHCMC05 vectors using the method described by Hardy (50). Transformants were spread onto BHI plates supplemented with 10 μg/mL of chloramphenicol and incubated at 37°C for 16-18 hours.

### Cloning and expression of *ddl6* and *ddl7*

The coding sequence of the genes for Ddl6 and Ddl7 (1032-bp) were amplified and cloned into the *Bam*HI and *Xho*I sites of pET28a vector (Novagen) using the primers TddlF and Tddl28aR (Table S4). The recombinant plasmids were transformed into *E. coli* BL21 (DE3) cells (New England Biolabs) and transformants were selected on LB agar supplemented with kanamycin (30 μg/mL). 1L LB was inoculated with 10 ml overnight culture, and expression was induced in exponential growth phase by addition of 0.5mM isopropyl β-D-1-thiogalactopyranoside for 3h at 37°C. Cells were harvested by centrifugation at 15,000xg at 4°C and stored at -20°C until protein purification.

### Purification of N- and C-terminal His-tagged Ddl6 and Ddl7 proteins by FPLC

Preserved cells were thawed on ice for 30 minutes and resuspended in buffer A (50 mM NaH2PO4, 300 mM NaCl, 10 mM imidazole, pH 7.4) containing cOmplete EDTA-free protein inhibitor cocktail (Roche, 1 tablet/10 ml), lysozyme (1.0 mg/ml) and DNAse (1 U/ μL). The suspension was incubated on ice for 30 minutes and sonicated using a sonicator (6 pulse, each 10 second). Cellular debris was removed by centrifugation at 15,000xg for 30 mins at 4°C. Recombinant both N-and C-terminally His-tagged proteins were purified using a Ni-NTA Superflow cartridge (1ml, Qiagen) and the Fast Protein Liquid Chromatography (FPLC) system (BioRad). The proteins were eluted with filter sterilized buffer B (50 mM NaH2PO4, 300 mM NaCl, 250 mM imidazole, pH 7.4). Protein concentration was determined using Pierce BCA Protein Assay Kit (ThermoFisher Scientific) and purity was checked by sodium-dodecyl sulphate polyacrylamide gel electrophoresis (SDS-PAGE). The gels were stained with Coomassie Blue (BioRad) and detained with destaining reagent (10% acetic acid and 20% methanol).

### Enzymatic activity assay

The D-ala-D-ala ligase activity of purified Ddl enzymes was assayed as described previously with some modifications (51, 52): *1) Formation of D-ala-D-ala dipeptide:* The reaction mixture containing the following components were incubated for 4 hours at 37°C: 50 mM Tris-HCl, 10 mM MgCl2, 10 mM KCl, 20 mM D-alanine, 2 mM ATP, 2.5 mM Glutathione and 20 μg/mL of purified Ddl. To stop the reaction, the mixtures were boiled for 5 min at 95°C. To determine the substrate specificity, an equimolar amount of D-Ser or D-Lac was substituted with D-Ala. Next, 4 μl of the reactions were loaded onto cellulose chromatography paper (Whatman) and ascending chromatography was developed for 3 hours in butanol-acetic acid-water (12:3:5 vol/vol/vol) containing 1.0% ninhydrin. The paper was dried for 5 minutes at 95°C. D-ala-D-ala dipeptide formation in the reaction catalysed by Ddl was identified by comparison with commercially available D-ala-D-ala (Sigma). *2) Measurement of inorganic phosphate in the Ddl-catalysed reaction:* The enzymatic activity of Ddl was also assayed by measuring the release of inorganic phosphate in the reaction (similar to the D-ala-D-ala dipeptide formation assay) with EnzChek phosphate assay kit (Life Technologies). 10 μl of the reaction product was added to 790 μl of standard reaction mixture containing 230 μl dH2O, 50 μl 20X reaction buffer, 200 μl 2-amino-6-mercapto-7-methylpurine riboside (MESG) substrate solution and 10 μl purine nucleoside phosphorylase and incubated at 22°C for 30 minutes. After incubation absorbance of the reactions was measured at 360 nm with a spectrophotometer (Ultrospec 2000, Pharmacia Biotech). The amount of inorganic phosphate released into the reactions was calculated from the standard curve of inorganic phosphate.

### Inhibition of Ddl activities by D-cycloserine

To observe the inhibitory activity of D-cycloserine on Ddl, different concentrations of the drug (5 mM to 80 mM) were added into the reaction catalysed by the enzymes (see above for *Measurement of inorganic phosphate in the Ddl-catalysed reaction*) and incubated for 4 hours at 37°C.

### Kinetic analysis of Ddl6 and Ddl7, DdlAEc and DdlTd

The V_max_ and K_m, D-ala2_ of Ddl6, Ddl7, DdlTd and DdlAEc were determined by using pyruvate kinase (PK)/lactate dehydrogenase (LDH)-coupled spectrophotometric system (52, 53).

All assays were conducted at 37° C in 96-well plate system using CLARIOstar® (BMG Labtech, Germany) in 200μL reaction buffer. The assay mixture (200μL) contained 100 mM HEPES (pH 7.5), 10 mM MgCl_2_, 10 mM KCl, 0.2 mM NADH, 6-10 U/mL PK, 9-14 U/mL LDH, 2 mM phosphoenolpyruvate (PEP) and variable concentrations of ATP and D-alanine (saturating concentrations were 500 μM and 100 mM for ATP and D-alanine, respectively). The reactions were initiated by adding Ddl enzymes at a final concentration of 0.0025 μg/μL. The initial velocity (v_0_) of the reactions was determined by measuring the changes in the absorbance at 340 nm for 500 secs (8.3 mins). The slope of the reaction progression curves (time (s) *vs* change in absorbance at 340 nm) is equivalent to the v_0_ (v_0_ = slope=?Y/?X)(54). The slope was calculated by the MARS data analysis software (BMG Labtech, Germany). After the v0 values for each substrate concentration (ATP and D-alanine) were obtained, GraphPad Prism version 7 (GraphPad Software, Inc., USA) was used to determine the V_max_ and K_m_ values by fitting the curves for Michaelis-Menten model.

The values for _Km,ATP_ of different Ddls were determined at a fixed saturating concentration of D-alanine (100 mM) and varying concentrations of ATP starting from 5 μM to 320 μM. The Km values for second D-alanine binding (K_m,D-ala2_) was determined by using a fixed saturating concentration of ATP (500 μM; approximately 10 times of K_m,ATP_). The values of K_cat_ (s^-1^) for all enzymes were determined by dividing the Vmax (μM/s) with the concentration of enzymes, [E] (μM).

### Phylogenetic analysis

The phylogenetic tree was constructed using MEGA6 software (55). The sequences of Ddl6 and Ddl7 and their homologs were aligned with Clustal Omega (55). The default settings for Neighbour-Joining method in MEGA6 were applied to construct the phylogenetic tree.

### Homology modelling of Ddl6 and ddl7

The homology models of Ddl6 and Ddl7 were constructed using i-Tasser server (http://zhanglab.ccmb.med.umich.edu/I-TASSER) (15). The amino acid sequences were submitted to the server and among the top five models built by the server, model 1 was selected based on the highest C-score and TM-score. A higher value of C-score signifies a model with a high confidence. A TM-score >0.5 indicates a model of correct topology and a TM-score <0.17 means a random similarity (56). Modelling was evaluated with PyMOL (The PyMOL Molecular Graphics System, Version 1.8 Schrödinger, LLC). The homology models of Ddl6 and Ddl7 were matched with each other with the TM-align structural alignment program (http://zhanglab.ccmb.med.umich.edu/TM-align) and the overlapped structure were evaluated with PyMOL.

### Determination of minimum inhibitory concentrations (MIC)

The MIC of D-cycloserine, vancomycin and beta-lactams were determined by agar dilution method following the guidelines of Clinical and Laboratory Standards Institute (CLSI). Three individual colonies of *E. coli* harbouring pET-*ddl6* or pET-*ddl7* and *B. subtilis* 168 strains harbouring pHCMC05-*ddl6* or pHCMC05-*ddl6* were inoculated in Mueller-Hinton (MH) broth supplemented with appropriate antibiotics and cultured at 37°C for 4-5 hrs and the OD_600_ was adjusted to 0.08 to 0.1 using sterile saline. MH agar plates supplemented with various concentration of antibiotics starting from 0.125 to 256 μg/ml were prepared and inoculated with the adjusted bacterial suspensions with a multipoint inoculator. The spots were allowed to dry for 30 minutes and the plates were then incubated aerobically at 37°C for 16-20 hrs. The MIC value was defined as the lowest concentration of the inhibitory compounds giving rise to no visible growth. MIC determinations were performed at least three times in triplicates. As a control, *E. coli* or *B. subtilis* 168 strains transformed with empty vectors were used.

### Molecular docking

The predicted 3D-structures of Ddl6 and its two single mutants, L164F and W259C, as well as Ddl encoded by house-keeping gene of *T. denticola* (DdlTd) were used as the targets for molecular docking analyses. The 3D structure of Ddl6 was applied in PyMol program to generate L164F and W259C mutants. These protein targets were used for molecular docking of six ligands including D-alanine, D-cycloserine, ATP, apigenin, quercetin and salvicine, after minimisation and equilibration using AMBER program. AutoDock SMINA (57) was used to determine the best binding pocket by exploring all probable binding cavities in the proteins. Then GOLD (Genetic Optimization for Ligand Docking) (58, 59) was used for docking the ligands into the SMINA-located binding site to perform flexible molecular docking analyses.

Genetic algorithm (GA) is used in GOLD ligand docking to thoroughly examine the ligand conformational flexibility along with partial flexibility of the protein (60). The maximum number of runs for the ligand was set to 20 and in each run a population size of 100 with 100,000 operations was employed. The number of islands was 5, and the niche size of 2 was considered). The default cut-off values of hydrogen bonds was set to 2.5Å (dH-X), and for the van-der-Waals distance it was 4.0Å. The GA docking was terminated when the top solutions attained the root mean square deviation (RMSD) values within 1.5 A °.

### Molecular dynamics (MD) simulations

After molecular docking, the best poses of D-cycloserine in the D-Ala binding site of wild-type and W259C mutant of Ddl6 were selected as the starting structures to run MD simulations for 50 ns. The MD simulations were carried out using the AMBER 16.0 software package (61). The force fields parameters for D-cycloserine were generated using the ANTECHAMBER module of the AMBER programme. Each system was solvated using an octahedral box of TIP3P water molecules. Periodic boundary conditions and particle-mesh Ewald (PME) method were employed in all the simulations (62). Particle-mesh Ewald method enabled us to calculate the ‘infinite’ electrostatics without truncating the parameters. During each simulation, all bonds in which the hydrogen atom was present were considered fixed, and all other bonds were constrained to their equilibrium values by applying the SHAKE algorithm (63). A cutoff radius of 12Å was used for the systems. Minimization was performed in two phases, and each phase was performed in two stages. In the first phase, ions and all water molecules were minimized for 1500 cycles of steepest descent followed by 1500 cycles of conjugate gradient minimization. Afterwards, the systems were minimized for a total of 5000 cycles without restraint wherein 2500 cycles of steepest descent were followed by 2500 cycles of conjugate gradient minimization. After minimizations, the systems were heated for 1000 ps while the temperature was raised from 0 to 300 K, and then equilibration was performed without a restraint for 1000 ps while the temperature was kept at 300 K. Sampling of reasonable configurations was conducted by running a 50 ns simulation with a 2 fs time step at 300 K and 1 atm pressure. A constant temperature was maintained by applying the Langevin algorithm while the pressure was controlled by the isotropic position scaling protocol used in AMBER 16 package program (61).

### Statistical Analysis

Statistical analysis was performed by using GraphPad Prism (GraphPad Software, Inc., USA; version 7.0). Statistical significance was calculated using one-way ANOVA. *P* values of ≤0.05 were considered statistically significant.

### GenBank accession numbers

The sequences of the 2024 bp PCR amplicons containing *ddl6* and *ddl7* were submitted separately to GenBank with the accession number of KU886208 and KU886209, respectively. The 4421 bp sequence of PCR product containing the upstream sequence of *intI* along with the downstream *ddl7* was submitted to GenBank with the accession number KY039278.

## Acknowledgments

We gratefully acknowledge the Commonwealth Scholarship Commission in the UK for supporting MAR (BDCA 2013-4) and Public Health England for supporting SJ through a research grant to KMR (award code JGALASR). We are thankful to Dr Haitham Hussain (UCL) for providing us with the pHCMC05 vector.

## Author Contributions

MAR; Conceptualization, experimental, analysis, manuscript writing and editing. FK; experimental, analysis, manuscript editing. SJ; experimental, analysis, manuscript editing. KMR; experimental, analysis, manuscript editing. PM; analysis, manuscript editing. APR; conceptualization, analysis, manuscript writing and editing.

## References

1. K. Bush et al., Tackling antibiotic resistance. Nat Rev Micro 9, 894–896 (2011).

2. V. M. D’Costa, K. M. McGrann, D. W. Hughes, G. D. Wright, Sampling the antibiotic resistome. Science 311, 374–377 (2006).

3. A. H. Holmes et al., Understanding the mechanisms and drivers of antimicrobial resistance. Lancet 387, 176–187 (2016).

4. C. M. Collis, R. M. Hall, Expression of antibiotic resistance genes in the integrated cassettes of integrons. Antimicrobial agents and chemotherapy 39, 155–162 (1995).

5. C. M. Collis, G. Grammaticopoulos, J. Briton, H. W. Stokes, R. M. Hall, Site-specific insertion of gene cassettes into integrons. Molecular microbiology 9, 41–52 (1993).

6. G. D. Recchia, R. M. Hall, Gene cassettes: a new class of mobile element. Microbiology 141 (Pt 12), 3015–3027 (1995).

7. G. Cambray, A. M. Guerout, D. Mazel, Integrons. Annual review of genetics 44, 141–166 (2010).

8. S. R. Partridge, G. Tsafnat, E. Coiera, J. R. Iredell, Gene cassettes and cassette arrays in mobile resistance integrons. FEMS Microbiol Rev 33, 757–784 (2009).

9. M. R. Gillings, Integrons: past, present, and future. Microbiol Mol Biol Rev 78, 257–277 (2014).

10. N. Coleman, S. Tetu, N. Wilson, A. Holmes, An unusual integron in Treponema denticola. Microbiology 150, 3524–3526 (2004).

11. F. C. Neuhaus, The enzymatic synthesis of D-alanyl-D-alanine. Biochemical and Biophysical Research Communications 3, 401–405 (1960).

12. I. Tytgat et al., DD-ligases as a potential target for antibiotics: past, present and future. Curr Med Chem 16, 2566–2580 (2009).

13. É. Guerin et al., The SOS Response Controls Integron Recombination. Science 324, 1034–1034 (2009).

14. Z. Feng, R. G. Barletta, Roles of Mycobacterium smegmatis D-alanine:D-alanine ligase and D-alanine racemase in the mechanisms of action of and resistance to the peptidoglycan inhibitor D-cycloserine. Antimicrobial agents and chemotherapy 47, 283–291 (2003).

15. J. Yang, Y. Zhang, I-TASSER server: new development for protein structure and function predictions. Nucleic acids research 43, W174–181 (2015).

16. Y. Zhang, J. Skolnick, TM-align: a protein structure alignment algorithm based on the TM-score. Nucleic acids research 33, 2302–2309 (2005).

17. D. Meziane-Cherif, F. A. Saul, A. Haouz, P. Courvalin, Structural and functional characterization of VanG D-Ala:D-Ser ligase associated with vancomycin resistance in Enterococcus faecalis. The Journal of biological chemistry 287, 37583–37592 (2012).

18. C. Fan, P. C. Moews, C. T. Walsh, J. R. Knox, Vancomycin resistance: structure of D-alanine:D-alanine ligase at 2.3 A resolution. Science 266, 439–443 (1994).

19. D. I. Roper, T. Huyton, A. Vagin, G. Dodson, The molecular basis of vancomycin resistance in clinically relevant Enterococci: Crystal structure of d-alanyl-d-lactate ligase (VanA). Proceedings of the National Academy of Sciences of the United States of America 97, 8921–8925 (2000).

20. S. Liu et al., Allosteric inhibition of Staphylococcus aureus d-alanine:d-alanine ligase revealed by crystallographic studies. Proceedings of the National Academy of Sciences of the United States of America 103, 15178–15183 (2006).

21. D. Wu et al., D-Alanine:D-alanine ligase as a new target for the flavonoids quercetin and apigenin. Int J Antimicrob Agents 32, 421–426 (2008).

22. U. N. Dwivedi et al., Treponema pallidum putative novel drug target identification and validation: rethinking syphilis therapeutics with plant-derived terpenoids. OMICS 19, 104–114 (2015).

23. C. M. Collis, R. M. Hall, Expression of antibiotic resistance genes in the integrated cassettes of integrons. Antimicrobial agents and chemotherapy 39, 155–162 (1995).

24. S. R. Partridge et al., Definition of the attI1 site of class 1 integrons. Microbiology 146 (Pt 11), 2855–2864 (2000).

25. O. Svartstrom, M. Mushtaq, M. Pringle, B. Segerman, Genome-wide relatedness of Treponema pedis, from gingiva and necrotic skin lesions of pigs, with the human oral pathogen Treponema denticola. PloS one 8, e71281 (2013).

26. S. G. Dashper, C. A. Seers, K. H. Tan, E. C. Reynolds, Virulence Factors of the Oral Spirochete Treponema denticola. Journal of dental research 90, 691–703 (2011).

27. S. C. Holt, J. L. Ebersole, Porphyromonas gingivalis, Treponema denticola, and Tannerella forsythia: the ‘red complex’, a prototype polybacterial pathogenic consortium in periodontitis. Periodontology 2000 38, 72–122 (2005).

28. B. Y. Wang, B. Chi, H. K. Kuramitsu, Genetic exchange between Treponema denticola and Streptococcus gordonii in biofilms. Oral Microbiology and Immunology 17, 108–112 (2002).

29. S. Tansirichaiya, M. A. Rahman, A. Antepowicz, P. Mullany, A. P. Roberts, Detection of Novel Integrons in the Metagenome of Human Saliva. PloS one 11, e0157605 (2016).

30. Y. W. Wu, M. Rho, T. G. Doak, Y. Ye, Oral spirochetes implicated in dental diseases are widespread in normal human subjects and carry extremely diverse integron gene cassettes. Applied and environmental microbiology 78, 5288–5296 (2012).

31. Y. W. Wu, T. G. Doak, Y. Ye, The gain and loss of chromosomal integron systems in the Treponema species. BMC Evol Biol 13, 16 (2013).

32. L. E. Zawadzke, T. D. Bugg, C. T. Walsh, Existence of two D-alanine:D-alanine ligases in Escherichia coli: cloning and sequencing of the ddlA gene and purification and characterization of the DdlA and DdlB enzymes. Biochemistry 30, 1673–1682 (1991).

33. T. J. Treangen, E. P. C. Rocha, Horizontal Transfer, Not Duplication, Drives the Expansion of Protein Families in Prokaryotes. PLOS Genetics 7, e1001284 (2011).

34. N. E. Caceres et al., Overexpression of the D-alanine racemase gene confers resistance to D-cycloserine in Mycobacterium smegmatis. Journal of bacteriology 179, 5046–5055 (1997).

35. M. Merker et al., Whole Genome Sequencing Reveals Complex Evolution Patterns of Multidrug-Resistant Mycobacterium tuberculosis Beijing Strains in Patients. PloS one 8, e82551 (2013).

36. C. A. Desjardins et al., Genomic and functional analyses of Mycobacterium tuberculosis strains implicate ald in D-cycloserine resistance. Nat Genet 48, 544–551 (2016).

37. R. Curtiss, L. J. Charamella, C. M. Berg, P. E. Harris, Kinetic and Genetic Analyses of d-Cycloserine Inhibition and Resistance in Escherichia coli. Journal of bacteriology 90, 1238–1250 (1965).

38. G. Baisa, N. J. Stabo, R. A. Welch, Characterization of Escherichia coli d-Cycloserine Transport and Resistant Mutants. Journal of bacteriology 195, 1389–1399 (2013).

39. P. C. Karakousis, Antimicrobial Drug Resistance. V. S. Georgiev, Ed., Infectious Disease (Humana Press, New York, 2009), vol. 1.

40. K. J. Forsberg et al., The shared antibiotic resistome of soil bacteria and human pathogens. Science 337, 1107–1111 (2012).

41. M. O. Sommer, G. Dantas, G. M. Church, Functional characterization of the antibiotic resistance reservoir in the human microflora. Science 325, 1128–1131 (2009).

42. K. J. Forsberg et al., Bacterial phylogeny structures soil resistomes across habitats. Nature 509, 612–616 (2014).

43. A. M. Moore et al., Pediatric fecal microbiota harbor diverse and novel antibiotic resistance genes. PloS one 8, e78822 (2013).

44. D. Whitford, Proteins: structure and function (J. Wiley & Sons, 2005).

45. C. S. Sevier, C. A. Kaiser, Formation and transfer of disulphide bonds in living cells. Nat Rev Mol Cell Biol 3, 836–847 (2002).

46. R. Sanchez et al., Protein structure modeling for structural genomics. Nature structural biology 7 Suppl, 986–990 (2000).

47. M. Arthur et al., Evidence for in vivo incorporation of D-lactate into peptidoglycan precursors of vancomycin-resistant enterococci. Antimicrobial agents and chemotherapy 36, 867–869 (1992).

48. I. S. Park, C. T. Walsh, D-Alanyl-D-lactate and D-alanyl-D-alanine synthesis by D-alanyl-D-alanine ligase from vancomycin-resistant Leuconostoc mesenteroides. Effects of a phenylalanine 261 to tyrosine mutation. The Journal of biological chemistry 272, 9210–9214 (1997).

49. M. M. Cowan, Plant Products as Antimicrobial Agents. Clinical Microbiology Reviews 12, 564–582 (1999).

50. K. G. Hardy, Bacillus cloning methods. In DNA Cloning D. M. Glover, Ed. (IRL Press., Oxford 1985), vol. 2.

51. J. B. Bruning, A. C. Murillo, O. Chacon, R. G. Barletta, J. C. Sacchettini, Structure of the Mycobacterium tuberculosis D-alanine:D-alanine ligase, a target of the antituberculosis drug D-cycloserine. Antimicrobial agents and chemotherapy 55, 291–301 (2011).

52. E. Daub, L. E. Zawadzke, D. Botstein, C. T. Walsh, Isolation, cloning, and sequencing of the Salmonella typhimurium ddlA gene with purification and characterization of its product, D-alanine:D-alanine ligase (ADP forming). Biochemistry 27, 3701–3708 (1988).

53. F. C. Neuhaus, The enzymatic synthesis of D-alanyl-D-alanine. I. Purification and properties of D-alanyl-D-alanine synthetase. The Journal of biological chemistry 237, 778–786 (1962).

54. H. B. Brooks et al., “Basics of Enzymatic Assays for HTS” in Assay Guidance Manual, G. S. Sittampalam et al., Eds. (Eli Lilly & Company and the National Center for Advancing Translational Sciences, Bethesda (MD), 2004).

55. K. Tamura, G. Stecher, D. Peterson, A. Filipski, S. Kumar, MEGA6: Molecular Evolutionary Genetics Analysis version 6.0. Mol Biol Evol 30, 2725–2729 (2013).

56. Y. Zhang, J. Skolnick, Scoring function for automated assessment of protein structure template quality. Proteins 57, 702–710 (2004).

57. D. R. Koes, M. P. Baumgartner, C. J. Camacho, Lessons learned in empirical scoring with smina from the CSAR 2011 benchmarking exercise. Journal of chemical information and modeling 53, 1893–1904 (2013).

58. G. Jones, P. Willett, R. C. Glen, A. R. Leach, R. Taylor, Development and validation of a genetic algorithm for flexible docking1. Journal of molecular biology 267, 727–748 (1997).

59. G. Jones, P. Willett, R. C. Glen, Molecular recognition of receptor sites using a genetic algorithm with a description of desolvation. Journal of molecular biology 245, 43–53 (1995).

60. J. W. Nissink et al., A new test set for validating predictions of protein-ligand interaction. Proteins 49, 457–471 (2002).

61. D. A. Case et al., The Amber biomolecular simulation programs. Journal of computational chemistry 26, 1668–1688 (2005).

62. T. Darden, D. York, L. Pedersen, Particle mesh Ewald: An N·log(N) method for Ewald sums in large systems. The Journal of Chemical Physics 98 (1998).

63. J.-P. Ryckaert, G. Ciccotti, H. J. C. Berendsen, Numerical integration of the cartesian equations of motion of a system with constraints: molecular dynamics of n-alkanes. Journal of Computational Physics 23 (1977).

