## Supplementary Figures and Tables for "Integron Gene Cassettes Harboring Novel Variants of D-Alanine-D-Alanine Ligase Confer High-level Resistance to D-Cycloserine"

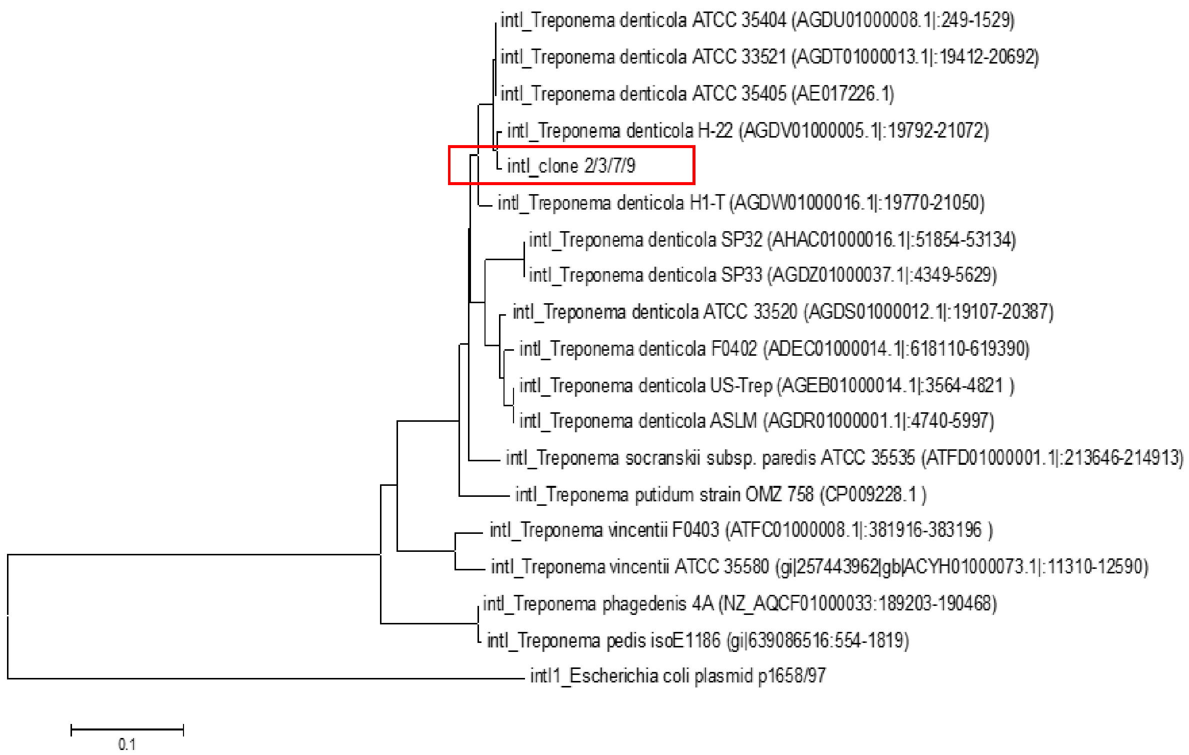


**Figure S1.** The phylogenetic tree of homologues of *intI* found upstream of *ddl7* in the 4421-bp inserts in pGEM-T Easy. The evolutionary relationship was inferred using Neighbour-Joining Method (1). Evolutionary analyses were conducted in MEGA6 (2). *intI1* of *E. coli* was used as outgroup.

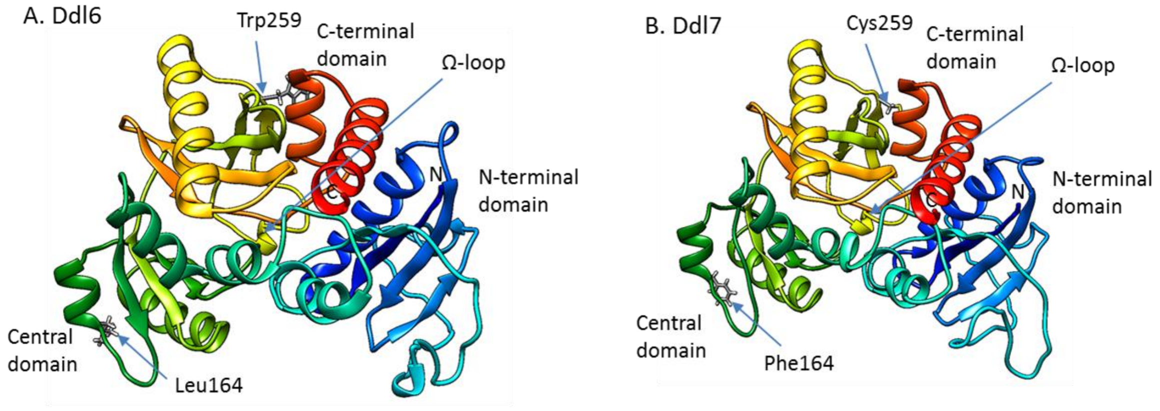


**Figure S2.** Predicted 3D-structures of Ddl6 (A) and Ddl7 (B) monomer as determined by I-TASSER. The model was viewed by UCSF Chimera. The residues at 164 and 259 are shown. The N-terminal domain and C-terminal domains were shown in blue and red colour, respectively.

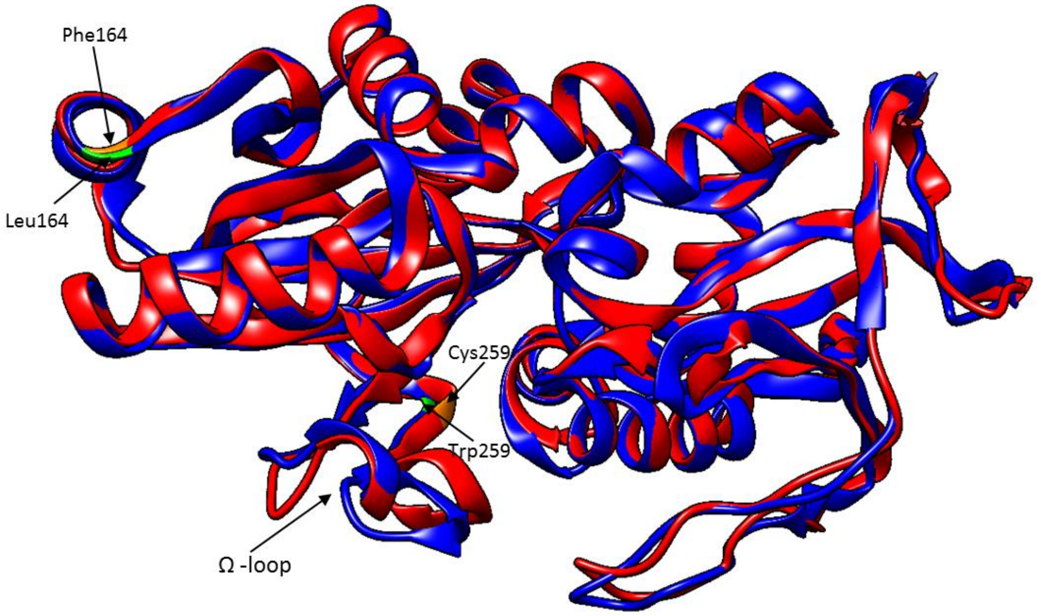


**Figure S3.** Superimposition of predicted 3D models of Ddl6 (blue) and Ddl7 (red). The alignment of the 3D structures of Ddl6 and Ddl7 were done using TM-align program (http://zhanglab.ccmb.med.umich.edu/TM-align/).

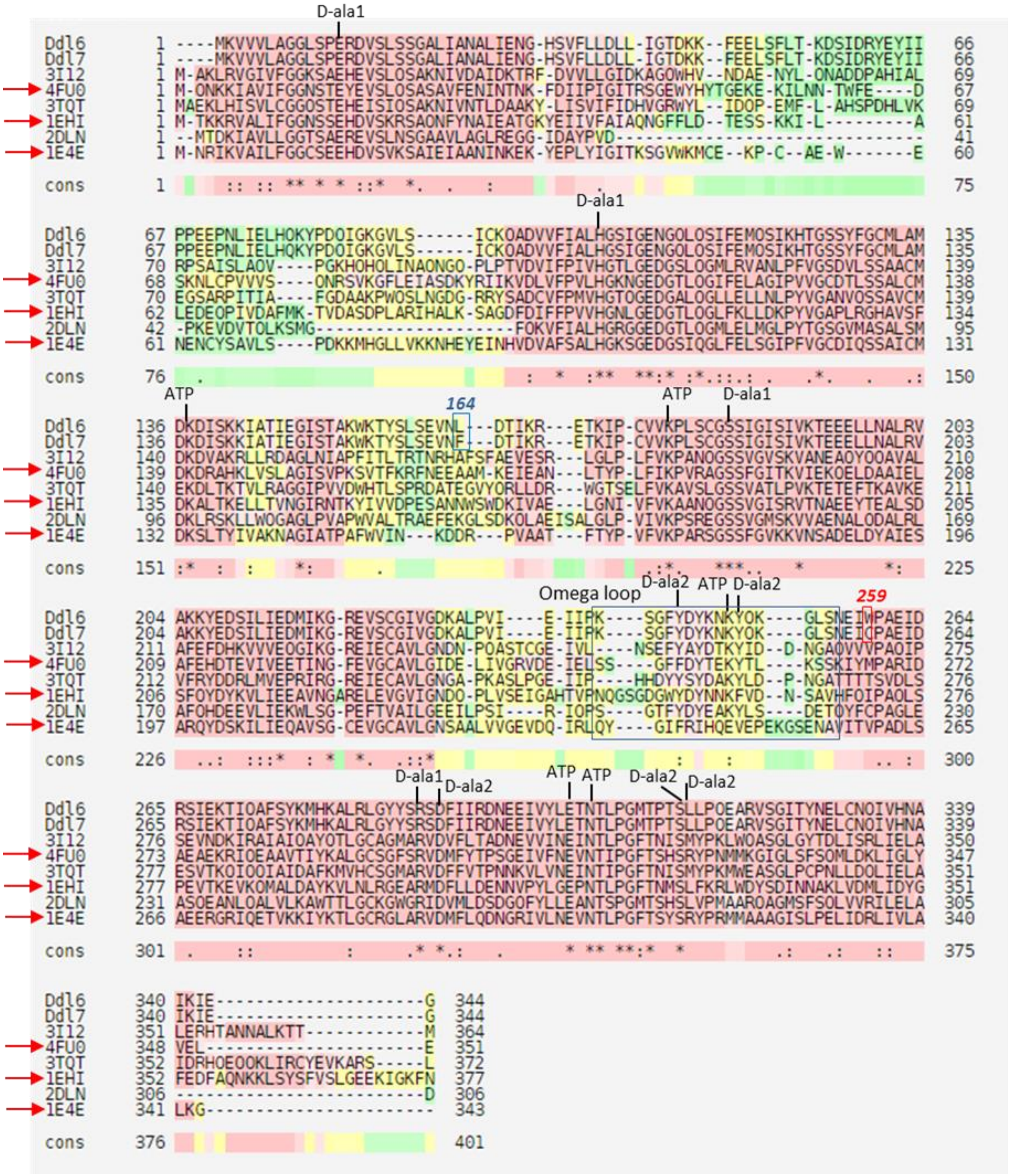


**Figure S4.** Multiple sequence alignment of the four proteins in PDB which have highest structural similarity to Ddl6 and Ddl7 (PDB codes: 3I12, 4FU0, 3TQT and 1EHI). The *E. coli* DdlB (PDB code: 2DLN) and *E. faecium* VanA (PDB code: 1E4E) were included in the alignment to compare the active/binding sites. PDB code 3I12: Ddl of *S. enterica* subsp. enterica Serover Typhimurium Str. LT2; 4FU0: D-ala-D-ser ligase (VanG) of E. faecalis; 3TQT: Ddl of *Coxiella burnetii*; 1EHI: D-ala-D-lac ligase of *L. mesenteroides* (LmDdl2). The vancomycin resistant proteins are marked with red arrows.

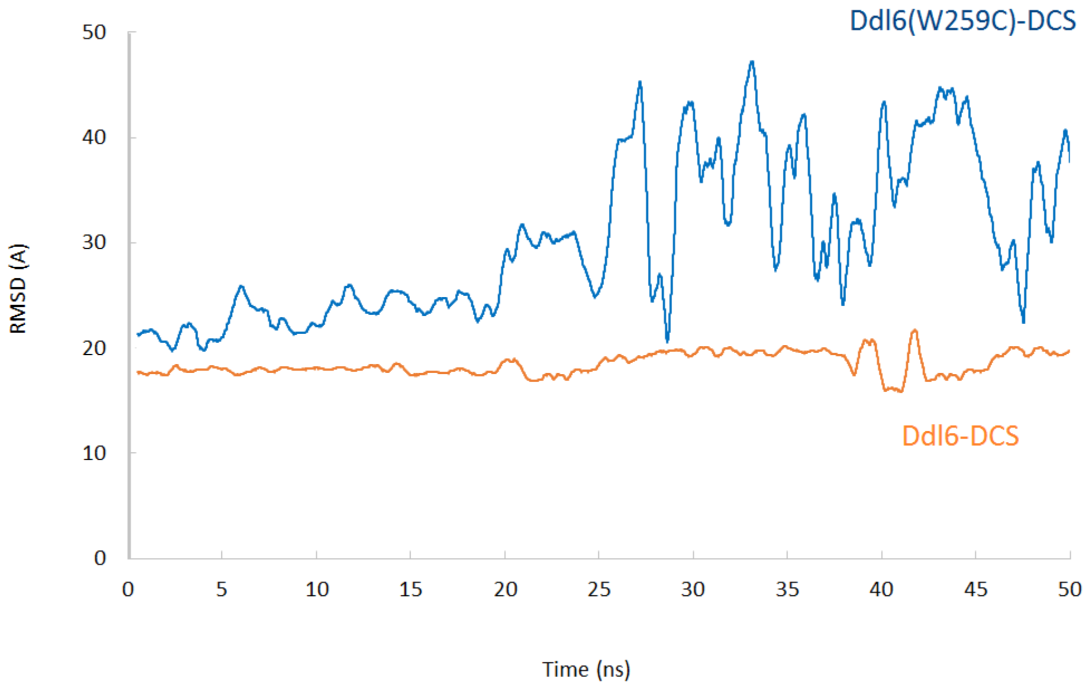


**Figure S5.** Monitoring the average movement of distance between the center of D-cycloserine and residue 259 (W259 and C259 of wild and mutant types respectively) in each complex during MD simulations.

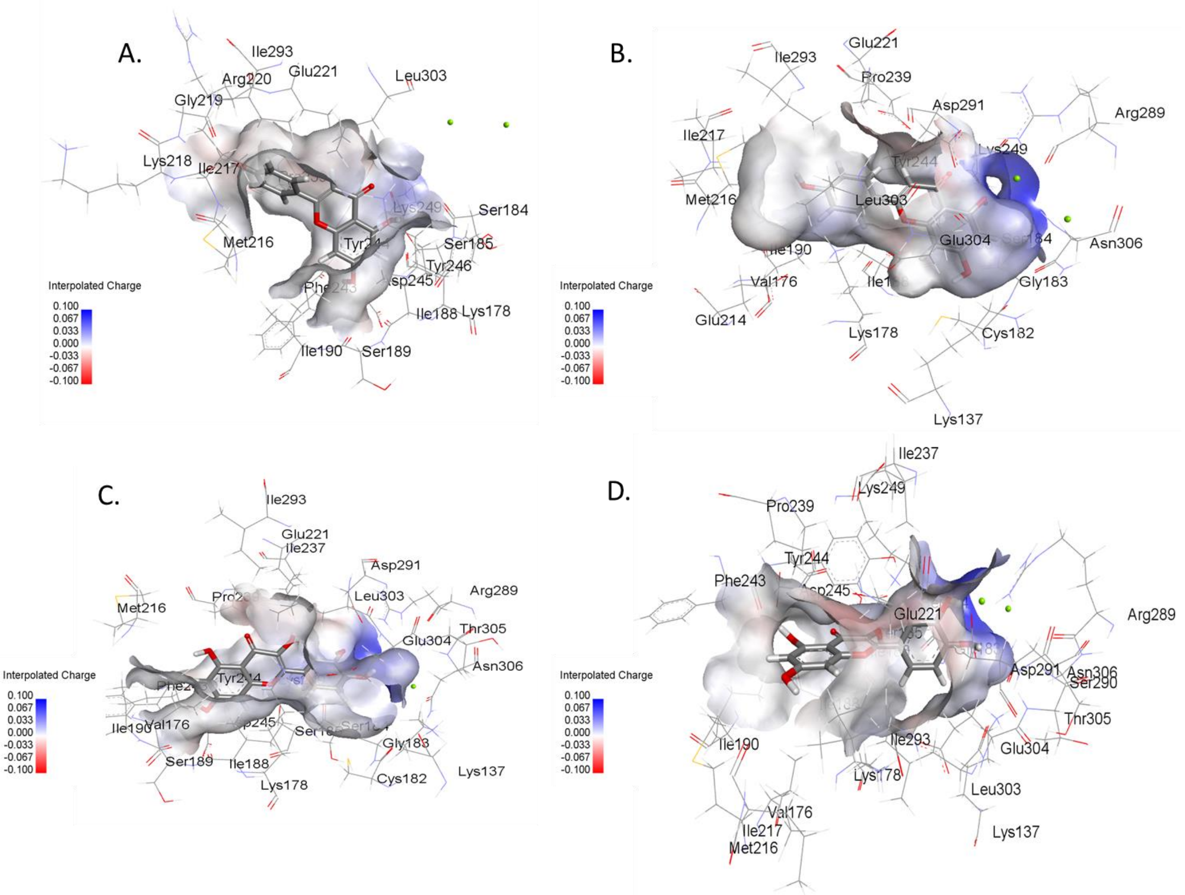


**Figure S6.** 3D structures of interaction of apigenin and quercetin with the ATP binding sites of WT Ddl6 and W259C mutant. A) Apigenin with WT Ddl6, B) Apigenin with W259C mutant, D) Quercetin with WT Ddl6 and D) Quercetin with W259C mutant.

**Table S1**. Top 10 structurally homologous protein of Ddl6 and Ddl7 in PDB. The I-TASSER models of Ddl6 and Ddl7 were used to compare the structural similarity with the PDB structures.

| Rank [a] | PDB hits | Resolution (Å) | Protein name, size and the host | TM-score | | RMSD[b] | | % identity [c] | |
|---|---|---|---|---|---|---|---|---|---|
| | | | | Ddl6 | Ddl7 | Ddl6 | Ddl7 | Ddl6 | Ddl7 |
| 1 | 3i12 | 2.2 | DdlA of *Salmonella enterica* subsp. *enterica* serovar Typhimurium str. LT2 (364 aa) | 0.943 | 0.966 | 1.82 | 1.20 | 28.44 | 28.74 |
| 2 | 4L1K | 2.3 | Ddl of *Xanthomonas oryzae* pv. Oryzae (384 aa) | 0.910 | 0.922 | 1.59 | 1.30 | 30.54 | 30.54 |
| 3 | 4FU0 | 2.35 | VanG (D-ala-D-ser ligase) of *Enterococcus faecalis* (351 aa) | 0.901 | 0.906 | 1.99 | 1.79 | 31.52 | 31.52 |
| 4 | 1EHI | 2.38 | D-ala-D-lac ligase (LmDdl2) of vancomycin-resistant *Leuconostoc mesenteroides* (377 aa) | 0.907 | 0.890 | 2.45 | 2.68 | 29.14 | 29.14 |
| 5 | 3TQT | 1.88 | Ddl of *Coxiella burnetii* | 0.886 | 0.596 | 1.89 | 1.71 | 27.63 | 27.63 |
| 6 | 1E4E | 2.5 | VanA (D-ala-D-lac ligase) of *E. faecium* BM4147 (343 aa) | 0.885 | 0.882 | 2.25 | 2.15 | 28.10 | 28.10 |
| 7 | 3SE7 | 3.07 | VanA, metagenomics (346 aa) | 0.872 | 0.876 | 2.02 | 1.81 | 26.43 | 26.43 |
| 8 | 2ZDH | 1.9 | Ddl of *Thermus thermophilus* HB8 (319 aa) | 0.867 | 0.859 | 1.91 | 2.09 | 31.53 | 31.53 |
| 9 | 2I8C | 2.46 | Ddl of *S. aureus* (358 aa) | 0.852 | 0.858 | 2.49 | 2.68 | 26.13 | 26.13 |
| 10 | 1IOW | 1.9 | Y216F mutant of DdlB of *E. coli* (306 aa) | 0.812 | 0.812 | 1.90 | 2.02 | 35.35 | 35.69 |

[a] Ranking of proteins is based on TM-score of the structural alignment between the query structure (Ddl6/Ddl7) and known structures in the PDB library;
[b] RMSD is the RMSD between residues that are structurally aligned by TM-align;
[c] % identity is the percentage sequence identity in the structurally aligned region

**Table S2.** Calculated energy contributions to form the Ddl6-D-cycloserine in wild and mutant complexes (kcal/mol) with standard errors of the mean (in parentheses); since the ligand was released from the receptor after around 20ns in mutant W259C form of Ddl6, so MMPBSA/MM-GBSA calculations were performed immediately before the ligand release.

| Energy distributions | Ddl6-D-cycloserine | Ddl6(W259C)-D-cycloserine |
|---|---|---|
| $\Delta E_{ele}$ | -28.63(4.31) | -18.68(3.73) |
| $\Delta E_{vdw}$ | -15.15(2.42) | -13.04(0.83) |
| $\Delta E_{sol}$ | 29.32(2.56) | 19.57(0.10) |
| $\Delta G_{PB}$ | -14.47(1.81) | -12.15(1.84) |
| $\Delta G_{GB}$ | -12.67(1.92) | -9.06(1.37) |

**Table S3.** Binding free energy (ΔG) and GOLD fitness score of apigenin, quercetin and salvicine for the D-alanine and ATP binding sites of native Ddl6 and its two single mutants.

| | **D-alanine binding site** | | | | | |
|---|---|---|---|---|---|---|
| | **Quercetin** | | **Apigenin** | | **Salvicine** | |
| | ΔG (kcal/mol) | Score | ΔG (kcal/mol) | Score | ΔG (kcal/mol) | Score |
| **Ddl6** | -40.20 | 33.42 | -43.41 | 35.48 | -43.81 | 41.76 |
| **L164F** | -39.90 | 32.33 | -44.61 | 37.21 | -44.87 | 41.95 |
| **W259C** | -39.85 | 32.95 | -41.58 | 33.02 | -44.93 | 41.58 |
| **DdlTd** | -40.20 | 33.43 | -39.90 | 32.34 | -40.14 | 39.13 |
| | **ATP binding site** | | | | | |
| **Ddl6** | -34.12 | 25.30 | -27.29 | 25.09 | -39.70 | 35.20 |
| **L164F** | -35.96 | 33.98 | -29.76 | 29.69 | -46.95 | 38.99 |
| **W259C** | -37.84 | 34.52 | -30.55 | 29.78 | -39.54 | 37.28 |
| **DdlTd** | -31.57 | 24.89 | -31.98 | 27.04 | -38.27 | 36.66 |

**Table S4.** Primers used in this study

| Name | Primer Sequence (5′-3′) | Target | Source |
|---|---|---|---|
| TDIF | TCAAGCCAAAATCAGGCTCT | *intI* in forward direction | (3) |
| MARS2 | GCAATGTCAGGTTGAAGC | *attC* in reverse direction | (3) |
| ddlF | GTAGTACTTGCTGGAGGATTA A | *ddl6/ddl7* (Forward primer) | This work |
| ddlR | GCCTTCAATTTTTATTGCATTA TGT | *ddl6/ddl7* (Reverse primer) | This work |
| TddlF | GCTA*GGATCC*ATGAAGGTAGT AGTACTTGC | *ddl6/ddl7* (Forward primer, *Bam*HI site underlined) | This work |
| TddlR | ACGC*TCTAGA*TTAGCCTTCAAT TTTTATTGC | *ddl6/ddl7* (Reverse primer, XbaI site underlined) | This work |
| Tddl28aR | GCGG*CTCGAG*TTAGCCTTCAA TTTTTATTGC | *ddl6/ddl7* (Reverse primer, XhoI site underlined) | This work |
| Ddl6-490F | TATCAGAAGTTAATTTTGATAC AATAAAGAGAGA | *ddl6* cloned into pGEM-T Easy (forward primer to change C of *ddl6* at c.490 to T) | This work |
| Ddl6-490R | ATGAATAAGTTTTCCATTTTGC TGTAG | *ddl6* cloned into pGEM-T Easy (reverse primer to couple with *Ddl*6-490F primer) | This work |
| Ddl6-777F | CAACGAAATATGTCCTGCTGA AATA | *ddl6* cloned into pGEM-T Easy (forward primer to change G of *ddl6* at c.777 to T) | This work |
| Ddl6-777R | GATAATCCTTTTTGATACTTAT TTTTATAATCATAAAAT | *ddl6* cloned into pGEM-T Easy (reverse primer to couple with *Ddl*6-777F primer) | This work |

| | | | |
|---|---|---|---|
| Upint-5100F | GGCGGAGATGAAGATACCCTT | 2.7 kb upstream of integrase of *T. denticola ATCC35404* | This work |
| EC-ddlA-F | GCTA*GGATCC*ATGGAAAAACT GCGGGTAGGAAT | *ddlA* of *E. coli* (forward primer, *Bam*HI site underlined) | This work |
| EC-ddlA-R | ACGC*AAGCTT*CATTGTGGTTTT CAATGCGTTATC | *ddlA* of *E. coli* (reverse primer, HindIII site underlined) | This work |
| DdlTD-F | GCGC*GAATTC*ATGAATATAGC AATCATTTACGGCGG | *ddl* of *T. denticola* (forward primer, *Eco*RI site underlined) | This work |
| DdlTD-R | ACGC*AAGCTT*AGATTGACGGC AGGTTTTGAGTTTTC | *ddl* of *T. denticola* (reverse primer, HindIII site Underlined) | This work |
| M13F | GTAAAACGACGGCCAG | M13 forward sequencing | Universal |
| M13R | CAGGAAACAGCTATGAC | M13 reverse sequencing | Universal |

**References:**


1. N. Saitou, M. Nei, The neighbor-joining method: a new method for reconstructing phylogenetic trees. *Mol Biol Evol* **4**, 406-425 (1987).
2. K. Tamura, G. Stecher, D. Peterson, A. Filipski, S. Kumar, MEGA6: Molecular Evolutionary Genetics Analysis version 6.0. *Mol Biol Evol* **30**, 2725-2729 (2013).
3. S. Tansirichaiya, M. A. Rahman, A. Antepowicz, P. Mullany, A. P. Roberts, Detection of Novel Integrons in the Metagenome of Human Saliva. *PloS one* **11**, e0157605 (2016).
